# Population coding of quantity discrimination in mouse posterior parietal cortex

**DOI:** 10.1101/2025.07.26.666982

**Authors:** Jikan Peng, Dong Yang, Chenshi Xu, Tian Xu

## Abstract

Quantity discrimination, the ability to identify preferable sizes or amounts of objects, is well-documented across humans and animals, yet its underlying neural mechanisms remain elusive. Here, we report that mice exhibit an innate quantity discrimination behavior for food amounts, consistent with the Weber-Fechner law. Inhibition of the Posterior Parietal Cortex (PPC), but not other cortex regions, abolishes this behavior. Two-photon calcium imaging reveals that the PPC neurons with preferential activities towards food amounts are detected at a given time; however, their preferential identities change over time. Conversely, the PPC exhibits population synchrony during quantity discrimination, and the levels of population synchrony correlate with food amounts. Together, our study identifies a central role of the PPC in quantity discrimination, providing a population coding for quantity discrimination.

## INTRODUCTION

Quantity discrimination is an essential ability for human and animal survival. This ability enables individuals to judge different sizes or amounts of objects for the preferred options. For example, human infants and chimpanzees choose a larger piece of cracker over a smaller one (*1, 2*). Similarly, when presented with prey of different sizes, cats and cuttlefish select the larger one (*3, 4*). Furthermore, when offered a choice between two sets of crackers with different amounts, human infants choose the set with more crackers (*1*). Such behavior is also widespread in non-human primates, canines, avian, amphibian, and even invertebrate species (*5–11*). In addition to food, quantity discrimination behavior is also important when judging conspecifics, predators, and other environmental cues (*12–18*). Therefore, quantity discrimination is crucial for animal survival and appears to be evolutionarily conserved. However, its underlying neural mechanisms, particularly in model organisms, are poorly understood.

Brain imaging studies have identified potential cortex regions associated with quantity discrimination in humans when individuals were subjected to judging different numbers of dots or numerical symbols (*19–30*). In addition to the sensory cortex for processing inputs, the Prefrontal Cortex (PFC) and the Posterior Parietal Cortex (PPC) were identified in these fMRI studies for quantity discrimination. In addition to brain imaging studies, electrophysiological recording in monkeys has also identified neuronal activity within the PFC, the PPC, as well as the Anterior Inferior Temporal Cortex (aITC), when animals were trained to recognize images with the same number of dots (*31–33*). Moreover, these electrophysiological studies identified a group of neurons in these three regions exhibiting activity preferentially tuned towards a given number of dots (*31–33*). Thereafter, it has been proposed that these specific neurons, or the “Number Neurons,” represent an abstract code for numbers (*34–36*). While these studies in primates have identified putative “Number Neurons” and implicated key brain regions, they largely rely on trained numerical tasks with abstract symbols. It therefore remains an open question whether similar neural coding principles underlie the innate, spontaneous quantity discrimination behaviors that are crucial for survival, such as choosing between different amounts of food.

Here, we have established a food-based assay to study spontaneous quantity discrimination in mice. Furthermore, using inhibitory chemogenetics, we have identified the functional involvement of the PPC. Finally, using in vivo two-photon calcium imaging, we have revealed that the PPC activity encodes quantity discrimination not through specialized “Number Neurons,” but through the changes in population synchrony levels. Our findings provide new insights into the neural basis of an evolutionarily conserved cognitive function.

## RESULTS

### A food-based assay reveals quantity discrimination behavior in mice

Whether mice share the evolutionarily conserved ability of innate quantity discrimination remains unknown. To address this, we developed a novel food-based assay in which individual naïve adult mice explored an open-field arena containing two piles of food in opposing corners (Fig. 1, A and B). We evaluated the animals’ foraging preference by measuring the accumulative time spent in the zone surrounding each pile over one hour. A representative foraging path illustrates a clear preference for the zone containing the larger food pile (Fig. 1C and Fig. S1). When presented with 20 g versus 80 g of food in the assay, significant disparities were observed for the times that individual mice spent in different foraging zones, resulting in measurements of 20.17 ± 2.13% (mean ± SEM) and 67.34 ± 2.80%, respectively (Fig. 1D). Moreover, the amounts of food consumed from the 80 g pile was approximately twice that from the 20 g pile (Fig. S2A). No such disparity was observed when equal amounts of food were presented in the assay (40 g versus 40 g; Fig. 1D and Fig. S2A). In addition to the C57BL/6J strain, the 129 and the BALB/c strains of mice exhibited similar behavior (Fig. 1E and Fig. S2B). Finally, the quantity discrimination behavior was also observed when additional different amounts of food were presented (Fig. 1F and Fig. S3; table S1). These results reveal that mice can distinguish different amounts of food and show a preference towards more food, exhibiting a spontaneous quantity discrimination behavior.

**Fig. 1.**
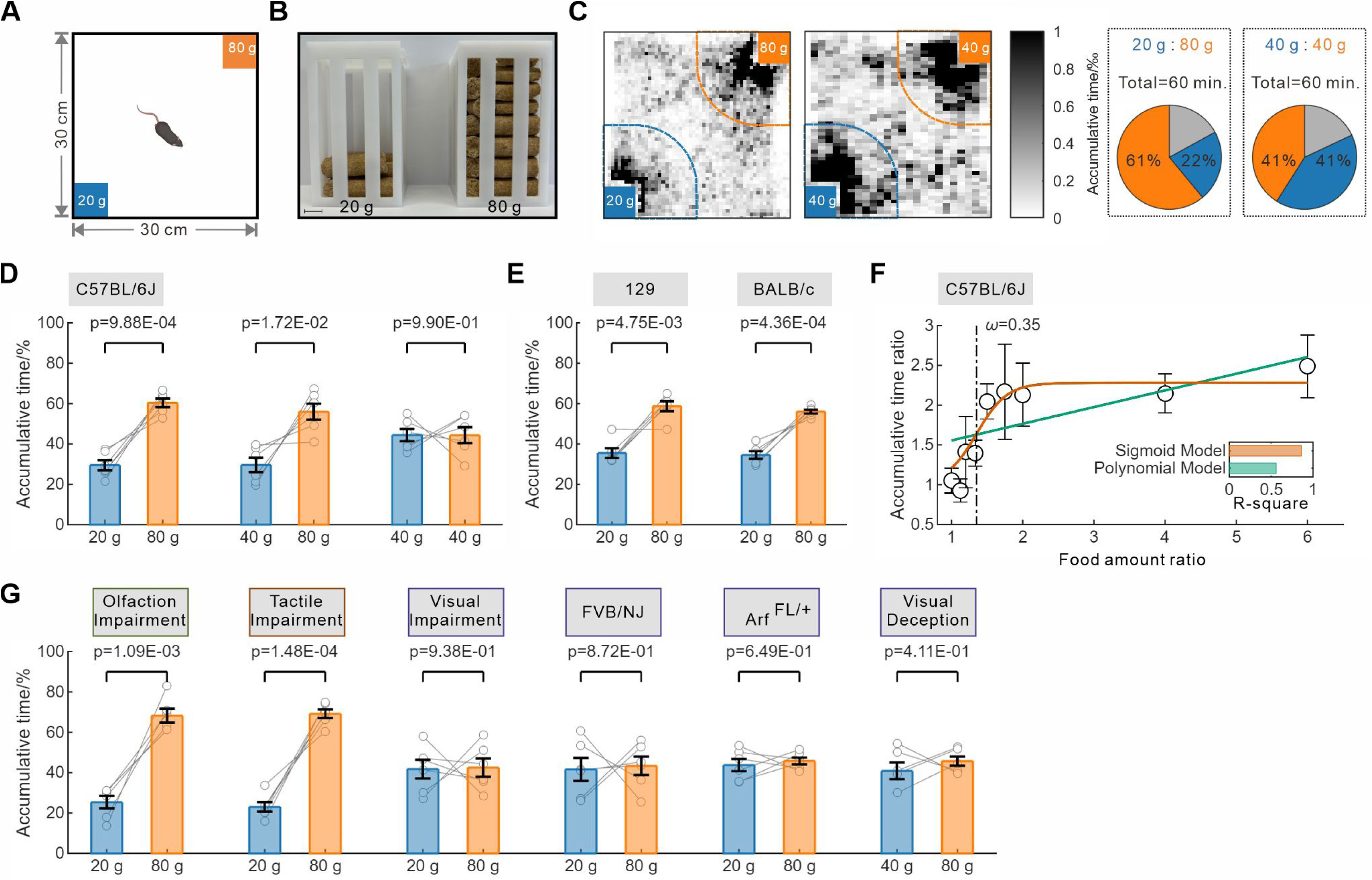
Quantity discrimination for food amounts in mice. (**A**) Schematic representation of different amounts of food in an open arena in a food-based quantity discrimination assay. (**B**) Representative food containers holding either 20 g or 80 g of food. Scale bar, 1 cm. (**C**) Tracking map illustrating the percentage of time the animal spent in each foraging zone (20 g versus 80 g, or 40 g versus 40 g). (**D**, **E**) Quantity discrimination behavior for different ratios of food amounts in the C57BL/6J (**D**), 129 and BALB/c (**E**) mice. (**F**) The relationship of the food amount ratio versus the accumulative time ratio fits with either the sigmoid (yellow line) or the polynomial (green line) model. The corresponding R-square values indicate model fitness, while the estimated Weber fraction (ω) is marked by the dotted line. (**G**) Quantity discrimination behavior following block of specific sensory inputs including chemical-induced olfactory anosmia, tactile sensory deprivation by whisker removal, or visual impairment due to crushing of optic nerves, retinal detachment, blindness, or visual deception of food amount. For **D**, **E** and **G**, two-tailed paired t-test; mean and SEM are shown; dots manifest data from individual mice (N=6 mice per group).

### Quantity discrimination in mice follows the Weber-Fechner law

In human quantity discrimination studies, the relationship between the different numbers or ratios of dots and the discriminating ability has been examined, which revealed that quantity discrimination in humans follows the Weber-Fechner law (*37*). The Weber-Fechner law is a general principle in behavioral science describing the relationship between the intensity of the physical stimuli and the perceived strength of these stimuli (*38, 39*). We therefore examined the relationship between different amounts or ratios of food and the quantity discrimination behavior in mice and confirmed that quantity discrimination in mice also follows the Weber-Fechner law (Fig. 1F, Fig. S2A, and Fig. S3; table S1). The Weber-Fechner law also specifies a threshold for discriminating distinct stimulus intensities, known as the Weber fraction (ω) indicating the strength of perception ability (*38, 39*). Our experiments revealed a Weber fraction value of 0.37 for food quantity discrimination in mice (Fig. 1F). Together, these results indicate that quantity discrimination in mice follows the Weber-Fechner law, paralleling the observations of quantity discrimination in humans.

### Visual input is required for food quantity discrimination in mice

We next sought to the sensory modality influencing the food quantity discrimination behavior in mice. We first assessed the contribution of olfactory sensation by inducing temporary anosmia with a chemical agent. Despite successful induction of olfactory anosmia (Fig. S4), the food quantity discrimination behavior remained intact (Fig. 1G and Fig. S2C). Subsequently, we evaluated the role of tactile sensation by removal of tactile bristles and found that the food quantity discrimination behavior maintained (Fig. 1G and Fig. S2C). In contrast, disruption of visual sensation via surgical ablation of retinal ganglion fibers resulted in a loss of the food quantity discrimination in mice (Fig. 1G and Fig. S2C). This observation was further confirmed by experiments with visually compromised mice, including FVB/NJ adults with retinal detachment and *Arf ^FL^*^/+^ transgenic animals with congenital blindness (Fig. 1G and Fig. S2C). To underscore the importance of visual cues in food quantity discrimination, we conducted a visual illusion experiment with wild-type animals, in which a plastic block was buried in food in the container with less food, visually aligning it with the container with more food (40 g with block, 80 g without block). Consistent with experiments of damaged visual system, visual illusion disrupted the food quantity discrimination (Fig. 1, D and G, and Fig. S2C). These findings reveal that visual input is required for food quantity discrimination in mice.

### The PPC is functionally required for quantity discrimination in mice

We subsequently investigated which brain regions might be functionally required for quantity discrimination in mice. As fMRI imaging and electrophysiological recording in humans and monkeys have revealed associated activities in several cortex regions with quantity discrimination (*19–24, 31–33*), we examined whether the PPC is functionally required for quantity discrimination for food in mice using inhibitory chemogenetics (Fig. S5). We found that inhibition of the PPC completely abolished the preference for the larger food pile (Fig. 2A and Fig. S2D). In contrast, inhibition of the mouse equivalent of aITC, the Postrhinal Cortex (POR), had no effect on this behavior (Fig. 2A and Fig. S2D). To discern if the disruption of the PPC activity might impact visual sensory function, we conducted a novel object recognition test and confirmed that visual sensory activity was unaffected by the PPC inhibition (Fig. S6; see Materials and Methods for more details). Together, our findings in mice provided the first evidence that the PPC is functionally required for quantity discrimination.

**Fig. 2.**
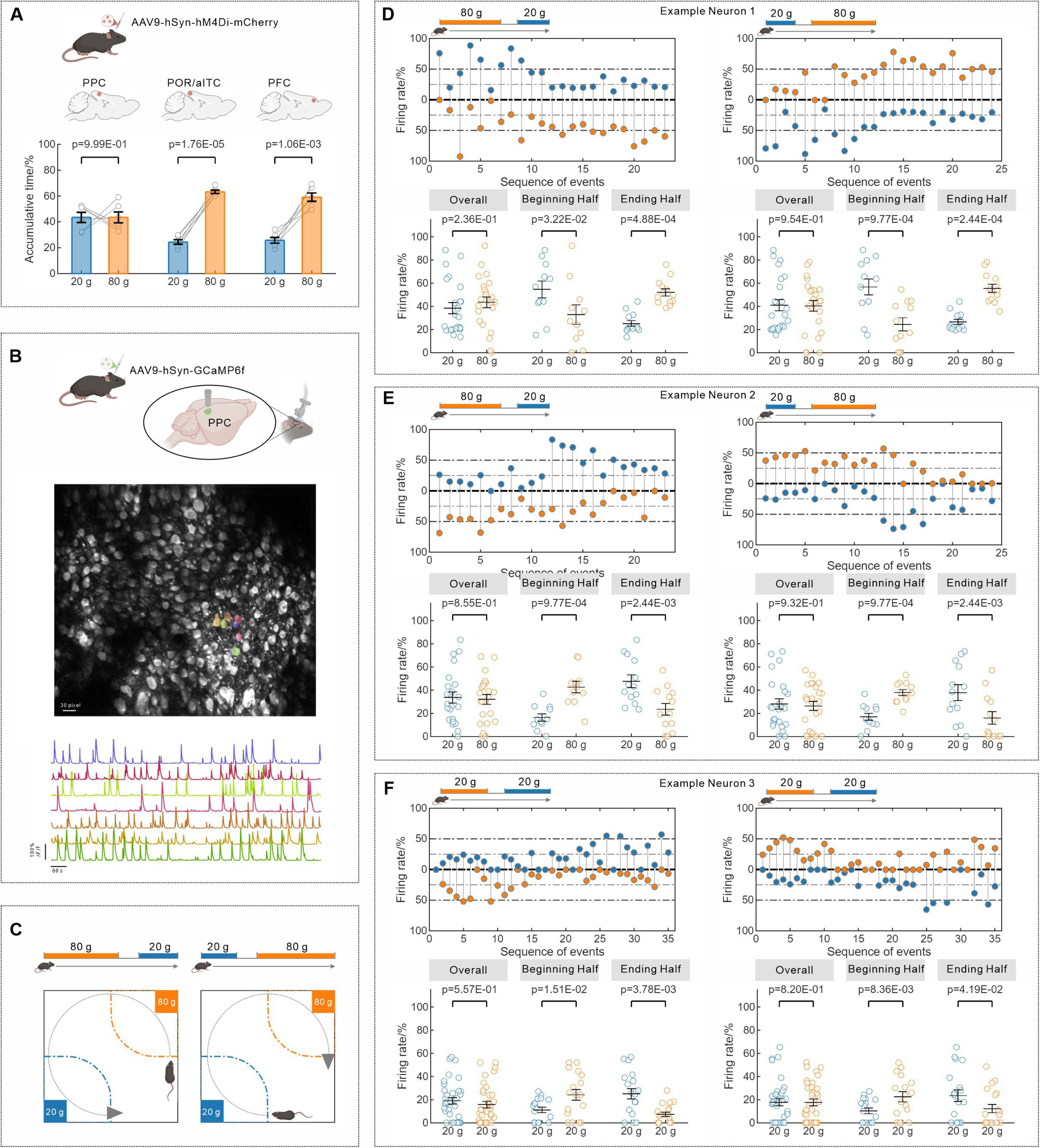
The involvement of the PPC for quantity discrimination. (**A**) Quantity discrimination behavior in mice following the inhibition of neuronal activity in the PPC, the POR/aITC, or the PFC. Two-tailed paired t-test; mean and SEM are shown; dots manifest data from individual mice (N=6 mice per group). (**B**) Two-photon imaging analysis for neuronal activity in layer II/III of the PPC during the quantity discrimination assay. Representative calcium signals from the PPC neurons are shown. (**C**) Two distinct events in which a mouse transitioned between two foraging zones. (**D**-**F**) Three representative neurons and their preferential activity towards different amounts of food over time. For each neuron, the upper panel shows neuronal activity for each behavior transition, and the lower panel tracks the preferential identities across different time periods. Two-tailed Wilcoxon signed-rank test; mean and SEM are shown; dots manifest data from individual events.

### “Number Neurons” are not required for innate quantity discrimination in mice

We then investigated how the PPC activity encodes different amounts of food and discriminates them. In the electrophysiological studies of monkeys, specific neurons with preferential activity for different number of dots, or the “Number Neurons,” were identified in three cortical regions: the PPC, the POR/aITC, and the PFC (*31–33*). However, our experiments revealed that inhibition of the POR/aITC did not disrupt innate quantity discrimination in mice (Fig. 2A). Among the three cortex regions, 30% of the recorded neurons in the PFC are the “Number Neurons”, which is much higher than 10% and 5% detected in the PPC and the POR/aITC, respectively (*32*). Therefore, to further investigate whether the “Number Neurons” are involved in innate quantity discrimination, we inhibited the PFC, which has the most “Number Neurons”, and found that this inhibition also did not disrupt food quantity discrimination behavior (Fig. 2A and Fig. S2D). These findings suggested that the “Number Neurons” may not be required for innate quantity discrimination in mice.

As the “Number Neurons” were also identified in the PPC (*32, 33*), we decided to conduct a detail examination of the neuronal activity in the PPC during quantity discrimination in mice using in vivo two-photon calcium imaging technology. This technology allows one to continuously monitor the activity of more than hundred neurons in each freely moving animal over extended periods (*40*). Briefly, we virally expressed the calcium indicator GCaMP6f in the right PPC and recorded neuronal activity during the food quantity discrimination assay using two-photon microscopy (Fig. 2B and Fig. S7A) (*41*). The two-photon calcium imaging data was processed following standard procedures (*42–44*), and the activity of 7,378 neurons across 48 mice were recorded (table S2; Fig. S8, and Fig. S9).

We analyzed the activity of individual neurons in response to 20 g versus 80 g of food, and did detect about 10% of the PPC neurons showing preferential activity towards either the 20 g or 80 g of food throughout the recording period (Fig. S10, A-C). This finding is very similar to the results from the electrophysiology studies in monkeys (*32, 33*). To further investigate such preferential activity, we defined two distinct behavioral events in which a mouse moved from one foraging zone to another (Fig. 2C and Fig. S7B). However, our analysis revealed that the PPC neurons had altered their preferential activity towards the different food amounts over time (Fig. 2, D and E, Fig. S10D). In other words, the “Number Neurons” had altered their preferential identities over time. Finally, we examined the “Number Neurons” phenomenon when the animals were presented with the equal amounts of food (20 g versus 20 g of food). We found that neurons with preferential activity were still detected at a given time and again such preferential identities changed over time (Fig. 2F and Fig. S10). Together, these findings indicate that “Number Neurons” are not required for innate quantity discrimination in mice.

### Population synchrony in the PPC is modulated by quantity discrimination

Since the unstable activity of individual neurons failed to explain quantity discrimination, we next asked whether quantity information is encoded at the population level in the PPC. First, we found that there is no significant difference in the overall firing magnitude when the animals were presented with different amounts of food (Fig. S11). We then discovered robust population synchrony in the PPC during quantity discrimination, with a Jaccard Index value as high as 0.7 (Fig. 3A; see Materials and Methods for more details). The average duration of the population synchrony is 1.95 ± 0.02 seconds (table S2 and Fig. S12). Furthermore, we discovered that all the recorded PPC neurons are activated during quantity discrimination and participated in population synchrony (Fig. S13). Together, these data revealed that the PPC exhibits population synchrony during quantity discrimination.

**Fig. 3.**
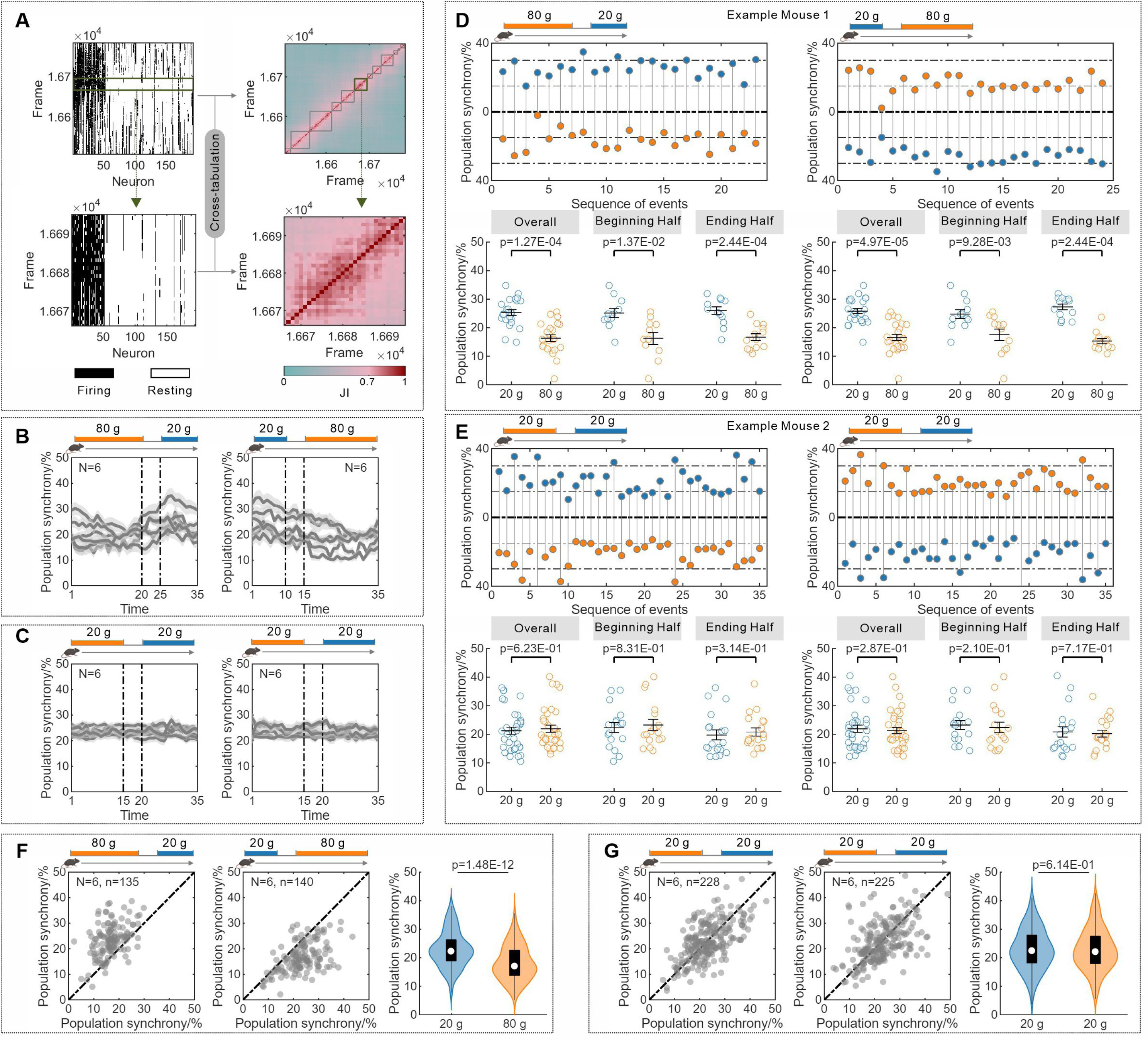
Population synchrony in the PPC during quantity discrimination. (**A**) The activation patterns of the PPC neuronal population at any given time (left), and the synchronous firing of a group of neurons (right, gray or green blocks), as indicated by the Jaccard Index (JI) calculation. (**B**, **C**) Changes in the level of population synchrony when the animal transitioned between foraging zones (**B**, 20 g versus 80 g; **C**, 20 g versus 20 g). Each line represents the mean ± SEM trajectory for a mouse (N=6 mice per group). (**D**, **E**) Two typical mice and their population synchrony levels over time. For each neuron, the upper panel shows population synchrony for each behavior transition, and the lower panel tracks the changes of population synchrony across different time periods. Two-tailed Wilcoxon signed-rank test; mean and SEM are shown; dots manifest data from individual events. (**F**, **G**) The levels of population synchrony in the PPC during quantity discrimination assay for 20 g versus 80 g of food (**F**), 20 g versus 20 g of food (**G**). Two-tailed Mann-Whitney U test; N indicates the number of mice; n indicates the number of events.

Critically, we discovered that the levels of population synchrony in the PPC are precisely modulated by different amounts of food. First, when the animals moved from the 20 g to the 80 g foraging zone, the levels of population synchrony decreased (Fig. 3B). Conversely, an opposite trend was observed when they transitioned from the 80 g to the 20 g foraging zone (Fig. 3B). No such differences were observed when equal amounts of food were presented in the assay (20 g versus 20 g; Fig. 3C). Second, unlike previous observations in “Number Neurons,” such changes in the levels of population synchrony persisted over time (Fig. 3, D and E). Finally, when the animals were presented with either 20 g or 80 g of food, the corresponding levels of population synchrony were 23.33 ± 0.76% and 16.61 ± 0.75%, respectively (Table 1; Fig. 3F, and Fig. S14). In contrast, when the animals were presented with the same amount of food (20 g versus 20 g), the level of population synchrony remained the same (Table 1; Fig. 3G, and Fig. S14). Furthermore, when additional different amounts of food were examined (40 g, 60 g, 70 g, and 90 g), the levels of population synchrony changed consistently (Table 1; Fig. 4A, Fig. S14 and S15). Together, these data reveal that population synchrony in the PPC is modulated by quantity discrimination.

**Fig. 4.**
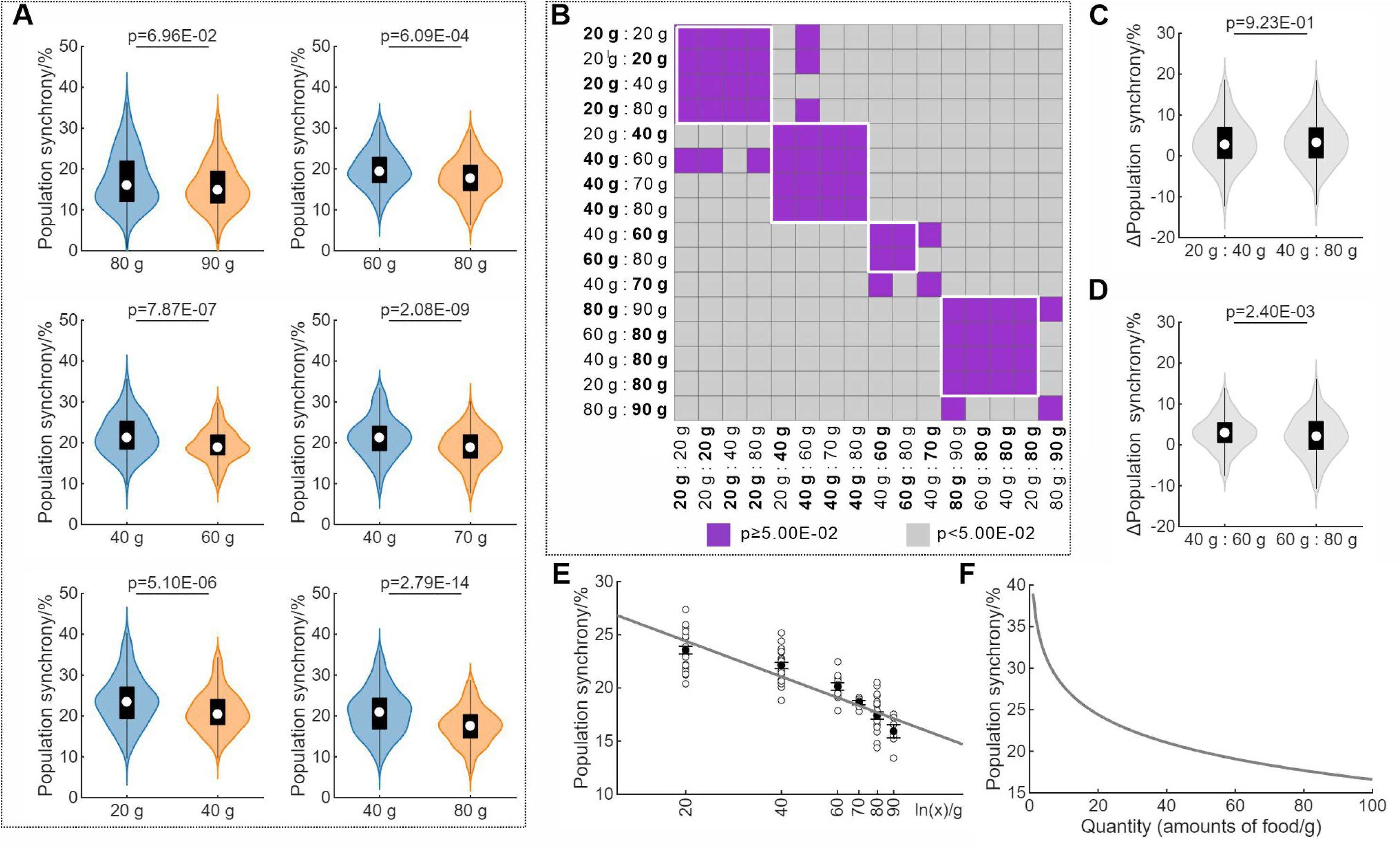
Population synchrony in the PPC correlates with quantity discrimination. (**A**) The levels of population synchrony in the PPC during quantity discrimination assay for different amounts or ratios of food (N=6 mice per group). (**B**) Heatmap of p-values from pairwise comparisons of population synchrony levels across different conditions. For clarity, the two conditions being compared in each instance are emboldened on their respective axes. (**C**) Relative differences in population synchrony levels when presented with 20 g versus 40 g of food, or 40 g versus 80 g of food. (**D**) Relative differences in population synchrony levels when presented with 40 g versus 60 g of food, or 60 g versus 80 g of food. (**E**) Relationship between population synchrony level and food amount fit to a logarithmic function (R-square=0.91). Mean and SEM are shown; dots manifest data from individual mice. (**F**) Predicted relationship between population synchrony level and food amount based on the logarithmic model. For **A**-**D**, statistical significance was determined by two-tailed Mann-Whitney U test.

**Table 1.** Levels of population synchrony during quantity discrimination.

| **Amounts of Food (g/g)** | **Levels of Population Synchrony (%) <sup>a</sup>** |
| --- | --- |
| 20:20 | 23.34 ± 0.55:23.51 ± 0.66 |
| 80:90 | 17.90 ± 0.84:15.91 ± 0.57 |
| 60:80 | 20.02 ± 0.28:17.48 ± 0.52 |
| 40:60 | 22.49 ± 0.29:19.25 ± 0.30 |
| 40:70 | 22.25 ± 0.36:18.61 ± 0.18 |
| 20:40 | 24.02 ± 0.83:21.61 ± 0.95 |
| 40:80 | 22.08 ± 0.35:17.50 ± 0.38 |
| 20:80 | 23.33 ± 0.76:16.61 ± 0.75 |

<sup>a</sup> Values are presented as mean ±SEM (N=6 mice per group).

### Population synchrony in the PPC correlates with quantity discrimination

We subsequently explored the relationship between population synchrony levels in the PPC and quantity discrimination. First, for a given amount of food (*e.g.*, 20 g), the corresponding population synchrony level remained consistent regardless of the alternative option (*e.g.*, 20 g versus 20 g, 20 g versus 40 g, or 20 g versus 80 g; Fig. 4B), indicating an absolute coding of quantity information. Second, the relative differences in population synchrony were consistent for identical ratios of food amounts (20 g versus 40 g, or 40 g versus 80 g; Fig. 4C). Conversely, a significantly larger difference in population synchrony was observed between 40 g and 60 g of food than between 60 g and 80 g of food (Fig. 4D), revealing a non-linear scaling of population synchrony. Finally, our analysis showed the relationship between the level of population synchrony and the amounts of food follows a logarithmic function (Fig. 4, E and F). Together, our findings indicate that the levels of population synchrony in the PPC correlate with quantity discrimination.

### Neural Entropy model reveals a population coding for quantity discrimination

It is intriguing how the levels of population synchrony in the PPC could code the quantity information. The fundamental elements of population synchrony are the firing or the resting status of individual neurons, which is very similar to the on or the off status of the transistors on the computer chip (*45*). The quantity information or other contents coded by the binary statuses of the transistors on a chip is measured by Shannon’s Entropy (Fig. 5A; Materials and Methods) (*46*). By analogy, we proposed a Neural Entropy model to calculate the levels of population synchrony in the PPC or in any given population of neurons (Fig. 5B; Materials and Methods). Specifically, the firing or the resting status of each neuron behaves as a binary coding unit. The aggregation of pairwise synchronous firing for all neurons in the population or the level of population synchrony could be calculated accordingly following the Neural Entropy model (Fig. 5B; Materials and Methods). Using the recorded data of neuronal activities in the right PPC during quantity discrimination, we calculated the Neural Entropy values for the 20 g or 80 g of food quantity discrimination to be 0.14 ± 0.01 or 0.45 ± 0.01 bits, respectively, which is consistent with the quantity discrimination behavior (Fig. 5B). Furthermore, the Neural Entropy calculation based on the levels of population synchrony for the other different amounts of food also resulted in distinctive values consistent with the outcomes of the behavior (Fig. S16). Together, our Neural Entropy model indicates that population synchrony constitutes a population coding for quantity discrimination in the mouse PPC.

**Fig. 5.**
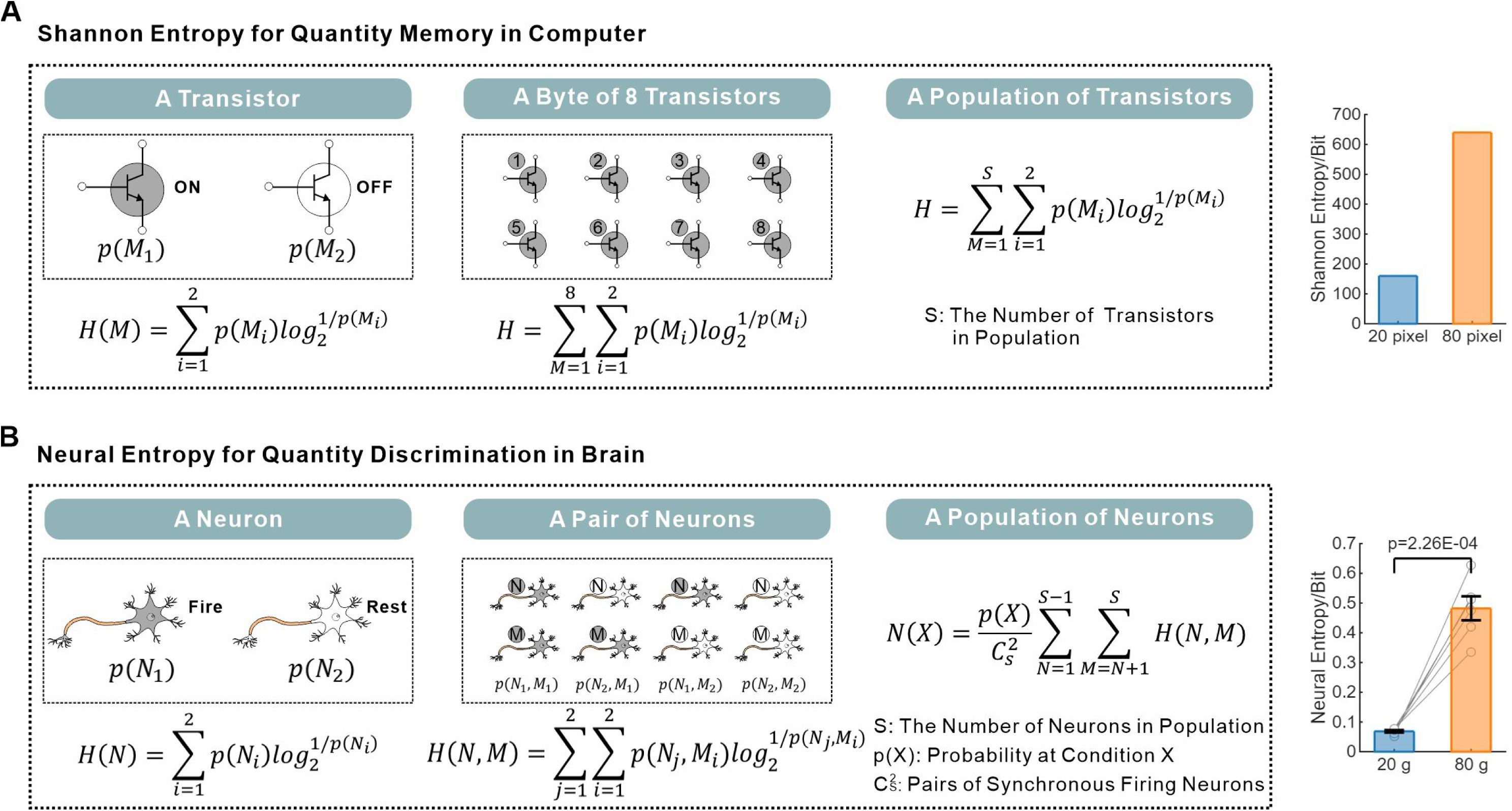
Neural Entropy model for quantity discrimination. (**A**) Calculation of Shannon Entropy for a single memory transistor, a Byte of eight transistors, and a population of transistors on a chip. The specified entropy values for encoding images with either 20 or 80 pixels. (**B**) Calculation of Information content for a single neuron, a pair of neurons, and a population of neurons in the PPC. The Neural Entropy values for population synchrony are calculated from the neuronal activity of the PPC during quantity discrimination for 20 g versus 80 g of food. Two-tailed paired t-test; mean and SEM are shown; dots manifest data from individual mice (N=6 mice per group).

## DISCUSSION

Quantity discrimination enables individuals to identify preferable sizes or amounts of objects, such as food, that are essential for animal survival. Here, we have established the first food-based assay to study quantity discrimination behavior in a model organism. Different strains of mice display an innate quantity discrimination ability for different amounts of food. Our analysis revealed that this ability for food amounts relies on the stimulus input from visual perception. Furthermore, the relationship between stimulus intensity of food amounts and quantity discrimination in mice follows the Weber-Fechner law, which is consistent with the results in humans for discriminating different numbers or dots (*37, 38, 47*). Compared to humans, monkeys, and crows, which have been used to study quantity discrimination or numeracy in the past, the mouse as a model organism offers many advantages for experimental manipulation and mechanistic exploration. Finally, different from discriminating dot numbers or sound beats in the past studies, quantity discrimination for different amounts of food is an innate behavior without training. Such an innate assay offers unique advantages in reducing the complexity and confounding factors introduced by learning and training.

Multiple cortex regions have been previously suggested to be involved in quantity discrimination through fMRI imaging and electrophysiological recording for discriminating dot numbers or numerical symbols in humans and monkeys (*19–33*). Using inhibitory chemogenetics, we have now shown that inhibition of the PPC, but not the PFC or the POR/aITC, blocks food quantity discrimination behavior in mice. Additionally, our companion study has shown that patients with surgery involved in the PPC fail in quantity discrimination and there is a genetic association between the PPC and intelligence for quantity discrimination (*48*). Overall, our findings have, for the first time, identified that the PPC is functionally required for quantity discrimination. Recent studies have shown that the PPC is also involved in navigation decision (*49–51*). Specifically, mice were trained for navigation decision in discriminative responses to images of dots, stripes, or different sound tones (*52–68*). Together, our work and the previous studies indicate that the PPC is responsible for discriminative decisions.

The electrophysiological recording in monkeys for recognizing the number of dots identified the existence of specific neurons exhibiting activity preferential towards specific numbers of dots, which led to the hypothesis that these “Number Neurons” represent an abstract code for numbers (*31–36*). However, multiple lines of evidence from this work and the navigation decision studies are against the existence of such “Number Neurons” and their role in innate quantity discrimination. It is important to note that the “Number Neuron” hypothesis was established based on studies in monkeys that were trained for numeracy tasks with abstract symbols. By contrast, our study investigates innate quantity discrimination for food amounts in naïve mice without training. The differences in the examined species and the experimental assays may account for the observed differences in mechanisms for quantity discrimination. First, our results showed that inhibition of the PFC or the POR/aITC, two regions containing “Number Neurons” (*31–33*), did not affect innate quantity discrimination in mice. Second, when the animals were presented with the equal amounts of food, neurons with different preferential activity were still detected. Third, and more importantly, our experiments revealed that the neurons with preferential activity alter their preferential identities over time. Similar results were also reported in the navigation decision studies, in which two-photon analysis revealed that the neurons in the PPC with preferential activity were detected at a given time and such preferential activity altered over time (*56, 59, 64*). Therefore, “Number Neurons” are not required for innate quantity discrimination. Furthermore, the “Number Neurons” coding hypothesis also has shortcomings in theoretical consideration for innate quantity discrimination or number. First, if it requires a special group of “Number Neurons” to code for each number, a vast number of neurons are needed to code for many numbers. By contrast, we and others have found that the levels of population synchrony or the states of the PPC neuronal population discriminates different quantity or navigation conditions. In other words, the PPC functions by determining differences in quantities or conditions in the given situations, which could be applied to numerous settings. The most unsettling issue is the “Number Neurons” coding hypothesis does not explain which number is bigger, which is essential to quantity discrimination in determining the preferred option(s) for animal survival. Together, both experimental evidence and theoretical considerations do not support the “Number Neurons” coding hypothesis for innate quantity discrimination.

Our investigation finally revealed that the neural basis of quantity discrimination does not depend on the number of firing neurons, the firing magnitude, or specialized “Number Neurons,” but rather on the levels of population synchrony within the PPC. Particularly, we have discovered that the levels of population synchrony in the PPC correlate with the quantity discrimination behavior. Therefore, we have identified a population coding for this fundamental cognitive function. The fundamental elements of population synchrony are the firing and resting binary statuses of individual neurons in a neuronal population. Binary coding systems have been utilized throughout human history, including the Chinese Bagua, Africa Ifa, Braille for visually impaired, and the transistors in computer chips (*45, 69*). The information content for the computer chip(s) is coded by the binary statues of the collection of the transistors and calculated by Shannon’s Entropy (*46*). In analogy, we have proposed a Neural Entropy model for calculating the information content of population synchrony and discovered that the Neural Entropy values of population synchrony could predict the quantity discrimination behavior. It is intriguing to consider whether similar coding system operates in other neuronal populations of the brain. Finally, it is fascinating to speculate whether forthcoming studies of the coding and energy strategies of the PPC could facilitate the design of future chips and computers.

## Acknowledgments

We thank Lijun Xiang and Hongyan Yang for technical assistance and the Xu lab members for discussion. JKP and CSX are supported by Westlake University Predoctoral Fellowship. TX is grateful to Yale and HHMI for more than two decades of support.

## Author contributions

Conceptualization, JKP and TX; Methodology, JKP; Investigation, JKP and CSX; Visualization, JKP and DY; Funding acquisition, TX; Project administration, TX; Supervision, TX; Writing – original draft, JKP and TX; Writing – review & editing, JKP and TX.

## Data and materials availability

All data are available in the main text or the supplementary materials. Codes for behavior analysis or two-photon imaging analysis are available from the authors on request.

## Supplementary Materials

Materials and Methods

Supplementary Text

Figs. S1 to S16

Tables S1 to S2

References (*70–88*)

## Materials and Methods

### Experimental model

All experimental procedures were approved by the laboratory animal resources center at Westlake University. The male mice at age of 6-8 weeks from C57BL/6J, 129, BALB/c, FVB/NJ, and *Arf ^FL^*^/+^ strains were used for the quantity discrimination assay. Only C57BL/6J mice were used for inhibitory chemogenetics, two-photon imaging experiments, olfactory test and novel object recognition test. Mice were housed under a 12-hour day/12-hour night cycle at 22℃-26℃ and were restrained from food for 48 hours before quantity discrimination experiments.

### The food-based assay

Each mouse was individually placed at the center of an open field arena measuring 30×30×50 cm (length × width × height) for a 10-minute habituation period to acclimate to the environment (Fig. 1A). Subsequently, designed amounts of food (Lab Mice Diet) were placed at random at diagonally opposite corners of the arena. The behavior of each mouse was recorded with an overhead camera for 60 minutes. The remaining food in each container was subsequently measured to determine consumption. Recorded videos were analyzed with the DeepLabCut software, which enabled precise tracking of mouse movement (*70*). Specific anatomical features, including the nose, ears and tail, were annotated for a detailed assessment of behavior. Using a customized MATLAB program, the distance of the mouse’s nose to each food container was computed and used to calculate the percentage of time spent in proximity to interpret foraging preferences. In a series of experiments, different food amount ratios were tested using specific combinations, as outlined in Table 1 and table S1. In a set of visual deception experiment, 40 g and 80 g of food ratio 1:2 was used; however, conspicuity was manipulated by concealing a plastic block within the container holding 40 g to simulate a similar visual volume as the 80 g container.

### Evaluation of Weber-Fechner law in the context of quantity discrimination

We designed various food amount ratios (table S1) and measured the ratios of the accumulated time mice spent in designated foraging zones. To analyze the relationship, we applied two mathematical models including polynomial regression and a sigmoid function. The data were most accurately described by a sigmoid curve (Fig. 1F). From the sigmoid curve, we derived the Weber fraction, a metric that quantifies the smallest detectable difference in stimulus intensity perceivable by the subject (*38*). In this context, it was adapted to measure the discernible differences in food amounts. The inflection point of the curve, occurring at approximately 1.37 (Weber fraction value 0.37), was critical for determining this value.

### Sensory input for discriminating different food amounts

Tactile deprivation was induced by carefully trimming the whiskers of the mice 10 minutes prior to testing (*71*). Transient anosmia in the mice was induced by administering 100 μl of 5% ZnSO4 into each naris 24 hours before conducting the assay or olfactory test (*72*). Optic nerve damage was induced by crushing the optic nerve 24 hours before testing (*73*), and a mutation in the Pde6b gene attributed to the early onset retinal degeneration specifically in the FVB/NJ mice (*74*). Transient expression of Arf in the Cdkn2a locus resulted in aberrant postnatal proliferation in the eye and subsequent blindness in the *Arf ^FL^*^/+^ mice (*75*).

### Olfactory test

The olfactory test for rodents comprises two distinct phases (*76*). During the first phase (P1), containers with swabs soaked in water, which served as nonsocial cues, were placed at opposite corners of the open field arena. Subsequently, in the second phase (P2), one of the water-soaked swabs was replaced with a swab permeated with urine, providing a social olfactory cue (Fig. S4). Mice were allowed to explore freely and detect olfactory cues for a period of 10 minutes during both P1 and P2. The movement of individual mice was analyzed similarly to that in the food quantity discrimination assay. To evaluate olfactory preference in the mice, a Preference Index (PI) was calculated by measuring their discrimination between the nonsocial and social olfactory stimuli. The PI is determined by the ratio of the difference to the sum of the accumulative time spent in the zones with social or nonsocial cues. A positive PI value indicated an individual mouse’s interest in social cues.

### Stereotaxic viral injection, inhibitory chemogenetics, and two-photon imaging

Mice were positioned on the stereoscopic positioning instrument after deep anesthesia induced by 1.5 to 2.5% isoflurane in oxygen (*77*). The skull’s fascia was removed with 4% phosphate-buffered saline following the incision of the mouse scalp. The bregma point served as the coordination origin for identifying the virus injection locations in the POR/aITC (AP: –3.00 mm; ML: ±4.3 mm: DV: 0.35 mm), or the PPC (AP: –2.00 mm; ML: ±1.6 mm: DV: 0.35 mm), or the PFC (AP: 2.43 mm; ML: ±0.4 mm: DV: 1.20 mm) (*50, 78, 79*).

For experiments involving inhibitory chemogenetics, a total of 200 nl AAV9-hSyn-hM4Di-mCherry virus was slowly injected at a rate of 50 nl/min into the bilateral regions of the specific cortex regions (*77*). The quantity discrimination behavior or the novel object recognition of individual mice was assessed 30 minutes after the intraperitoneal injection of 5.0 mg/kg Clozapine N-Oxide (CNO). Following the behavioral tests, the mice underwent intracardial perfusion with 0.9% NaCl for 30 seconds and 4% paraformaldehyde for 5 minutes. Individual brains were dissected, fixed with 4% PFA for 12 hours, and subsequently dehydrated with 30% sucrose for 24 hours. The brains were then coronally sliced into 20 μm sections using a cryostat (Fig. S5) (*80*).

For the two-photon imaging experiments, a total of 300 nl AAV9-hSyn-GCaMP6f virus was injected into the PPC in the right hemisphere (*77*). Post-virus injection, a small window of 300 μm to 400 μm in diameter was drilled just above the PPC and covered with a 1 mm-high slide. The mice were allowed a two-week recovery period before the two-photon imaging experiments. An imaging baseplate was positioned over the slide and secured with dental acrylic. Fast high-resolution miniature two-photon microscopy (FHIRM-TPM) was employed, and its holder was mounted onto the baseplate. Images were simultaneously acquired at a predetermined frame rate during the behavior tests.

### Novel object recognition test

The Novel Object Recognition test was conducted to examine the integrity of an animal’s visual sensory input (*81*), which consisted of three distinct phases (Fig. S6). In Phase 1 (P1), each mouse was allowed to freely explore an empty arena for 10 minutes to acclimate to the environment. In Phase 2 (P2), two identical objects (green cylinders) were placed at opposite corners of the arena, and the mice were given another 10 minutes to explore. In Phase 3 (P3), one of the familiar objects was replaced with a novel object (a yellow cube), and the mice were again provided 10 minutes to explore the arena. The analysis of the mice’s exploratory behavior and the calculation of the PI followed a protocol similar to that used in the olfactory test. A positive PI value during P3 indicated a preference for the novel object.

### Two-photon imaging processing

Images of neuronal activities in the PPC layer II/III were captured in freely moving mice using FHIRM-TPM microscopy at a resolution of 600×512 pixels (487.80×416.20 μm) and a frame rate of 10 Hz (Fig. 2B). The two-photon images were processed using established protocols, including motion correction, noise reduction, and segmentation of neuronal components (*42–44, 82*). For motion correction, the NoRMCorre algorithm was employed to compensate for rigid and non-rigid translational movement (*44*). Specifically, a reference frame with most activated neurons was manually selected from a subset of frames (one frame chosen from every 500, for a total of 60 frames). Each frame in the subset was aligned to this reference. The aligned frames were then averaged to create an initial reference image for the registration of the entire video. The mean projection images, both before and after registration, were illustrated (Fig. S8, A and B). Furthermore, the DeepInterpolation method, based on an encoder-decoder network with skip connections, was applied to reduce Poisson noise within individual frames (*82*). The model, originally trained on high-frequency (30 Hz) data, was fine-tuned using our 10 Hz dataset, resulting in effective noise reduction (Fig. S8, C and D).

Segmentation of neuronal components was initially performed using established morphological methods (*43*). Briefly, the two-photon images were divided into blocks of 100 frames (Fig. S8E). For each block, both maximum and average intensity projections were calculated (Fig. S8E). The difference between these projections was subjected to an adaptive threshold filter to generate a periodic mask containing multiple components (Fig. S8E). These periodic masks were subsequently refined using morphological operations, such as hole filling. Subsequently, we developed a new strategy to integrate these periodic masks and automatically identify individual neuronal components, as described in a separate study. We observed that pixels belonging to the same neuronal structure were more likely to be consistently grouped into the same component across different periodic masks, providing a basis for identifying all pixels within a given neuronal component. Specifically, for any given pair of pixels, the Jaccard Index (JI) was calculated as the ratio of the number of blocks in which both pixels were classified into the same component to the number of blocks in which either pixel was identified in any component (*83*). Pixels with a JI value of 0.5 or higher were grouped together, representing the spatial footprint of different regions of interest (ROI). Consequently, a temporal mask was generated (Fig. S8F). To address potential ROI overlap, which may arise from a single neuron or from distinct neurons sharing similar imaging fields, we calculated the percentage of pixel pairs with a JI greater than 0.5 between two ROIs. A threshold of 80% or higher was used to determine whether the two ROIs originated from the same neuron. Overlapping areas were then merged to generate a spatial mask (Fig. S8G). This spatial mask was further refined using the signal quality of individual ROIs to generate a soma mask (Fig. S8H). This process resulted in the identification of three distinct soma-like components, which were defined as individual neurons (Fig. S8I).

### Extraction of neuronal fluorescence signal

Similar to the processing of two-photon images, neuronal signal extraction also followed established protocols (*43, 84, 85*). Briefly, the initial fluorescence signal of any given ROI in the spatial mask was calculated by the mean intensity of all associated pixels in each frame (Fig. S9, A and B). The neuropil fluorescence was estimated by averaging the intensities of an equivalent number of surrounding pixels (Fig. S9, A and B). An ROI was selected as neuron if its fluorescence signal exceeded the 3-sigma level of its neuropil for at least three consecutive frames (Fig. S9B). The fluorescence signal was further refined by subtracting neuropil contamination and removing polynomial trend. Next, a baseline filter was applied to normalize the fluorescence signal (Fig. S9, C and D). This process resulted in the calcium signal, expressed as ΔF/F, where ΔF represents the fluorescence change from the baseline (Fig. S9E). In the calcium signal, positive or negative transients were identified by isolating segments of the signal that exceeded the baseline by more than two-sigma and persisted for a minimum duration of three consecutive frames (*84*). The sigma value was determined separately: for positive transients, using measurements below the baseline; for negative transients, using measurements above the baseline (*84, 86*). Neurons exhibiting a positive-to-negative transient ratio greater than ten were selected for further analysis. Finally, the event signal was extracted from the calcium signal (Fig. S9F), which was based on a frame rate of 100 ms and a decay constant of 860 ms (*85*).

### Quantity discrimination decoding analysis

We conducted a detailed analysis of various neuronal features to decode the quantity discrimination behavior in mice. These features were calculated for either individual neurons or neuronal population within the PPC as described below. Notably, neurons located at the edges of the image were excluded to ensure accurate decoding.

Event Definition: We identified two distinct behavioral events where a mouse transitioned between foraging zones (Fig. 2C). For each event, we measured and analyzed the time spent in each zone (Fig. S7B). Specifically, we assigned normalized time units to different behavioral states: 20 units for time spent in the zone containing 80 g of food, 10 units for time in the zone with 20 g of food, and 5 units for transitional periods between zones (Fig. 3B); 15 units for time spent in the zone containing 20 g of food, 15 units for time in the zone with 20 g of food, and 5 units for transitional periods between zones (Fig. 3C).

Neurons with preferential activities: We measured the activities of individual neurons when the animal was located in the foraging zone for the specific amount of food. Specifically, the average firing magnitude (FM) and the average firing rate (FR) per frame were assessed for each neuron. For animals presented with a specific amount of food, the neurons with preferential activities (FM or FR) were identified throughout the 60 min observation period (Fig. S10, A and B). To determine whether preferential activities changed over time, we tracked the identities of the individual neurons and their preferential activities across time and compared the data between the beginning and ending halves. In 34 out of the 48 animals, all neurons changed preferential activities. Overall, changed preferential activities were detected in 7,352 out of the 7,378 recorded neurons (Fig. 3, D and E, and Fig. S10, D and E).

Population synchrony: Among all the fired neurons in each animal presented with a given amount of food, the average percentage of the neurons fired in the same frame or the level of population synchrony was calculated (Table 1 and Fig. S14). For the duration of population synchrony, the Jaccard Index (JI) similarity value greater than 0.7 in all the consecutive frames or more than 82% the same neurons fired in all the consecutive frames were used for calculation (Fig. 3A, and Fig. S12) (*83*).

Similarity Index: To determine whether synchronous firing patterns involved the entire PPC neuronal population or specific subsets, we calculated a Similarity Index (SI) by comparing neuronal activity across different time frames (Fig. S13). For each neuron, we computed its firing rate during two distinct synchronous firing periods. The SI was derived by first taking the absolute difference between corresponding neuronal firing rates, then averaging these differences across all neurons. Lower SI values (approaching 0) indicate greater similarity in population activity patterns between the compared periods, suggesting consistent involvement of the same neuronal ensembles.

Pairwise comparison of populations synchrony: A cross-analysis of population synchrony levels was conducted between the conditions of different experimental groups (Fig. 4B). For each comparison, data from a specific condition in one group were statistically compared to data from a related condition in another group (*e.g.*, the first “20 g” condition in the **20 g** versus 20 g group was compared to the “20 g” condition in the **20 g** versus 80 g group).

### Statistical information

In our study, we employed various statistical methods to analyze the data, including the paired t-test, the unpaired t-test, the Wilcoxon signed-rank test and the Mann-Whitney U test, all conducted as two-tailed tests. P-values from the Mann-Whitney U test were adjusted using the Benjamini-Hochberg method. P-values were reported using scientific notation and rounded to two decimal places. In our data presentation, we utilized a variety of graphical representations to effectively convey our findings. Specifically, bar graphs were employed to illustrate mean values accompanied by Standard Error of the Mean (SEM) as error bars, with individual data points represented as dots for clarity (*e.g.*, Fig. 1D). Additionally, violin plots were utilized to display median values along with the interquartile range (25%-75%), providing insight into the distribution of the data. The data distribution was further visualized using a kernel density estimate curve (*e.g.*, Fig. 3F). Moreover, a specific graph was used to illustrate the distribution of the original data, presented through median values and interquartile ranges (*e.g.*, Fig. S13).

## Supplementary Text

### Neural Entropy

Claude Shannon developed Shannon Entropy (*H*) to calculate the information content (*46*). Given a variable *X*, with possible outcomes *x*_1_,…, *x_s_*, which occur with probability *p*(*x*_1_),…, *p*(*x_s_*), the Shannon Entropy of *X* is defined as:

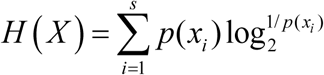

### Shannon Entropy for a single transistor

A transistor on a computer chip has on and off statuses, which store information content following Shannon Entropy, as given by:

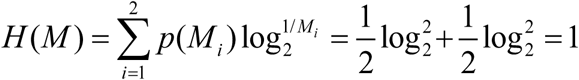

### Shannon Entropy for a population of transistors

A population of transistors on a computer chip constitutes a binary coding system (*87*), for which the information content can be expressed as the aggregate of the Shannon Entropy across S transistors in use, as calculated by:

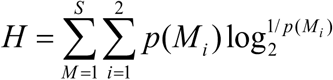

For instance, for encoding a 20-pixels or 80-pixels gray image, the corresponding Shannon Entropy value would be 160 bits or 640 bits, respectively (Fig. 5A).

### Information Entropy for a single neuron

A neuron also has binary conditions of fire or rest. Given a neuron (N), with its firing frequency *p*(*N*_1_) and its resting frequency *p*(*N*_2_), its information content could be similarly calculated by:

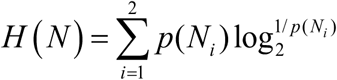

### Information Entropy for a pair of neurons

For a pair of neurons, they could fire synchronously or not (Fig. 5B). Joint Entropy is employed to quantify the information content for associated events (*88*). The Information Entropy for a pair of neurons ( *N*, *M*) could be calculated by:

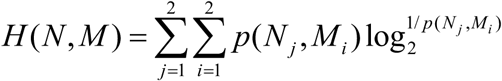

### Neural Entropy for a population of neurons

For a population of neurons or a brain region, it could code information by the levels of synchronous firing of the neurons. The information content could be calculated by the level of synchronous firing or pair-wise firing of the neurons in the population, termed as Neural Entropy (*N*). For a brain region performing a particular task ( *X*), it has a *S* number of neurons in its population and the pairs of synchronous firing neurons ( *C* ^2^). The information content could be calculated by measuring the level of synchronous firing or the average of the aggregated pair-wise firing, the Neural Entropy:

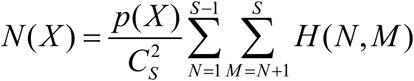

For instance, for presented with either a 20 g or 80 g of food, the corresponding Neural Entropy value would be 0.14 bits or 0.45 bits, respectively (Fig. 5B).

**Fig. S1.**
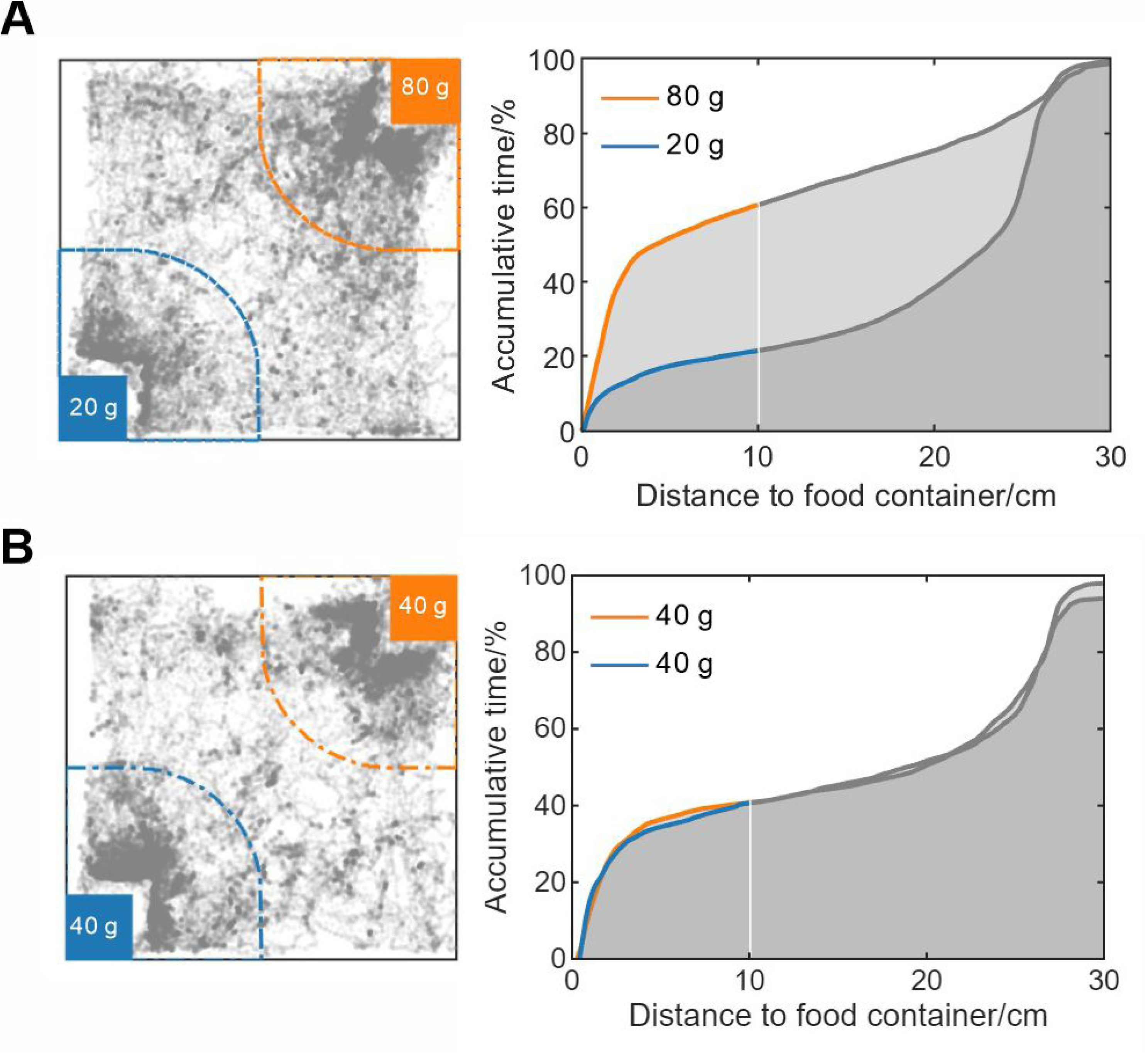
Typical tracking map in the food quantity discrimination assay. Representative tracking map illustrating the percentage of time the mouse spent in each foraging zone (**A**, 20 g versus 80 g; **B**, 40 g versus 40 g), along with its proximity to the food containers.

**Fig. S2.**
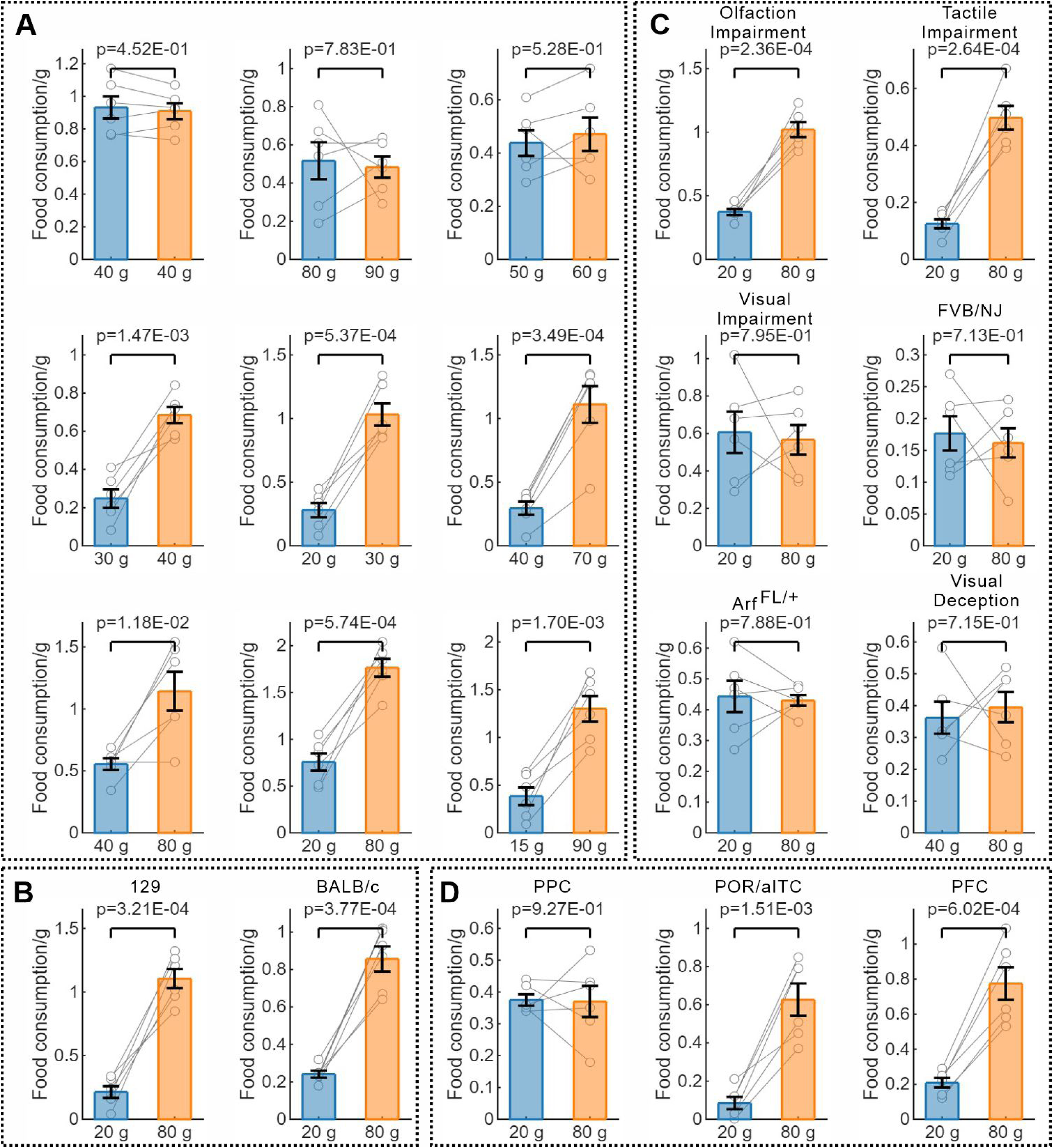
Food consumption in the food quantity discrimination assay. (**A**) Food consumption by C57BL/6J mice in quantity discrimination assays for different amounts of food. (**B**) Food consumption by 129 and BALB/c mice. (**C**) Food consumption by individual mice in sensory input deprivation experiments. (**D**) Food consumption by C57BL/6J mice in the experiments of inhibition of the PPC, the POR/aITC, or the PFC. For **A**-**D**, Two-tailed paired t-test; mean and SEM are shown; dots manifest data from individual mice (N=6 per group).

**Fig. S3.**
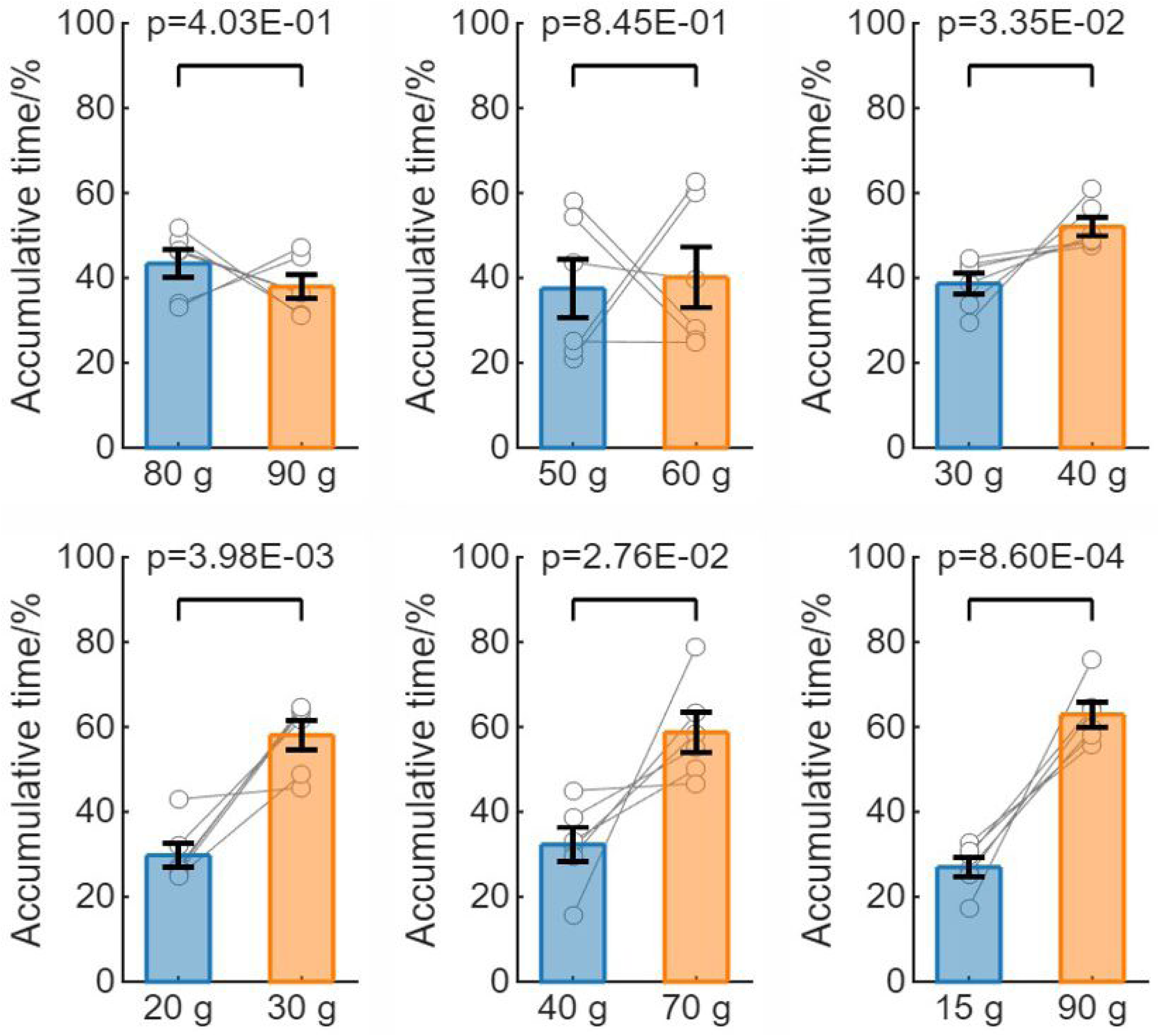
Quantity discrimination for additional food amounts in mice. Quantity discrimination behavior for different ratios of food amounts in the C57BL/6J. Two-tailed paired t-test; mean and SEM are shown; dots manifest data from individual mice (N=6 per group).

**Fig. S4.**
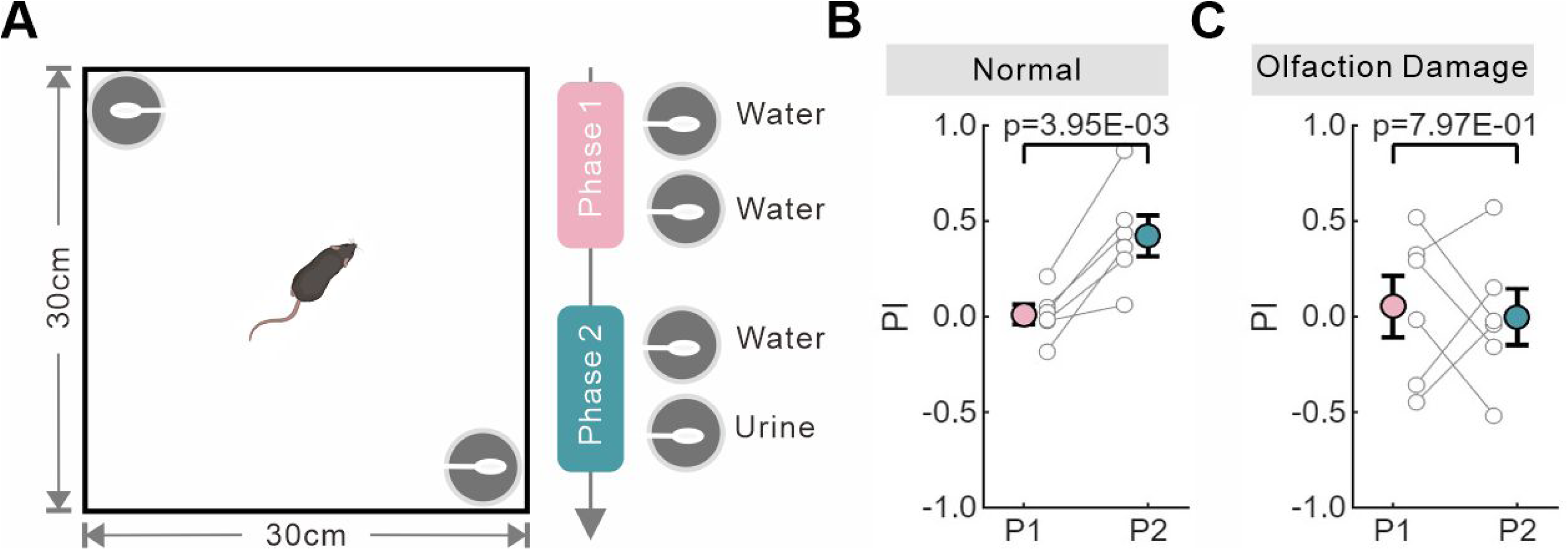
Olfactory test. (**A**) The exhibition of various olfactory cues in an open arena. (**B, C**) Preference Index (PI) values for olfactory cues during different experimental phases under normal condition (**B**) or following induced deprivation of olfactory sensation (**C**). Two-tailed paired t-test; mean and SEM are shown; dots manifest data from individual mice (N=6 per group).

**Fig. S5.**
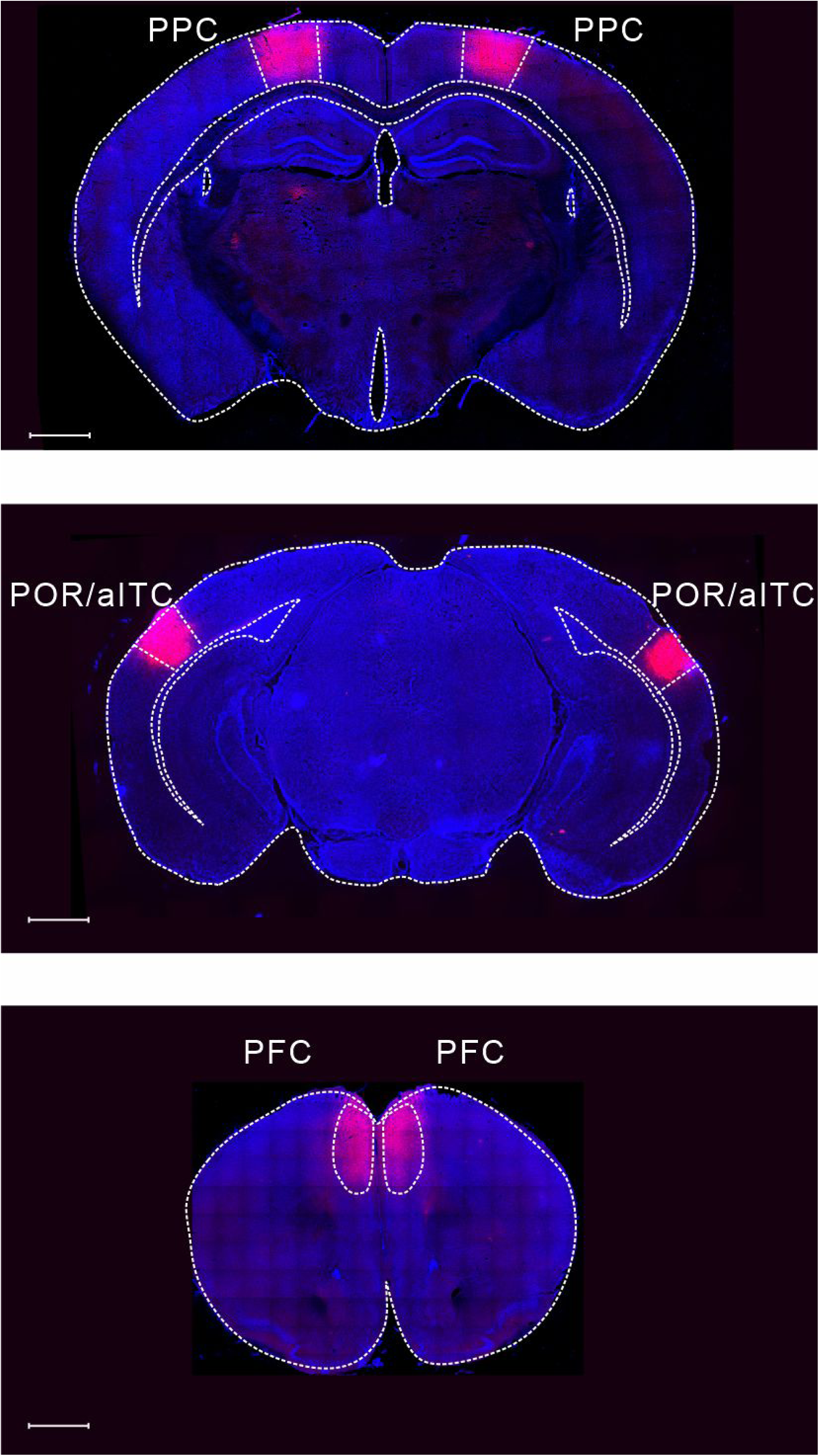
Inhibition of different cortex regions. Site-specific delivery of a viral vector expressing an inhibitory channel, AAV9-hSyn-hM4Di-mCherry, was performed in different cortex regions: the PPC (top), the POR/aITC (middle), or the PFC (bottom), as described in the Methods. Scale bars: 1 mm.

**Fig. S6.**
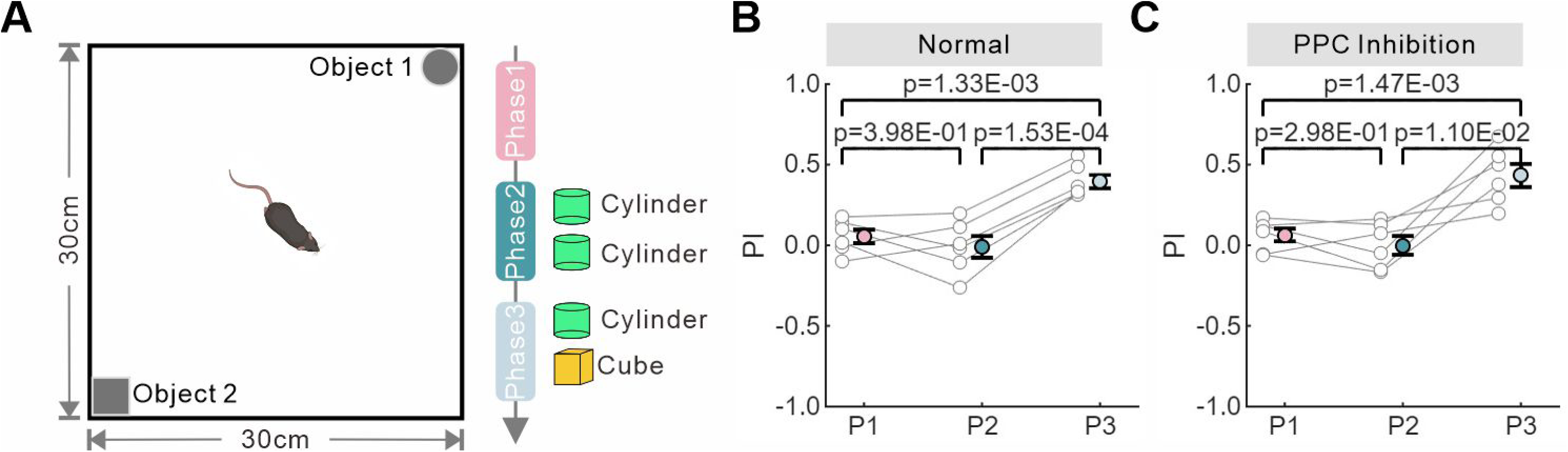
Novel object recognition test. (**A**) The positions of different objects in an open arena. (**B**, **C**) Preference Index (PI) values for objects during different experimental phases under normal condition (**B**) or following inhibition of neuronal activity in the PPC (**C**). Two-tailed paired t-test; mean and SEM are shown; dots manifest data from individual mice (N=6 per group).

**Fig. S7.**
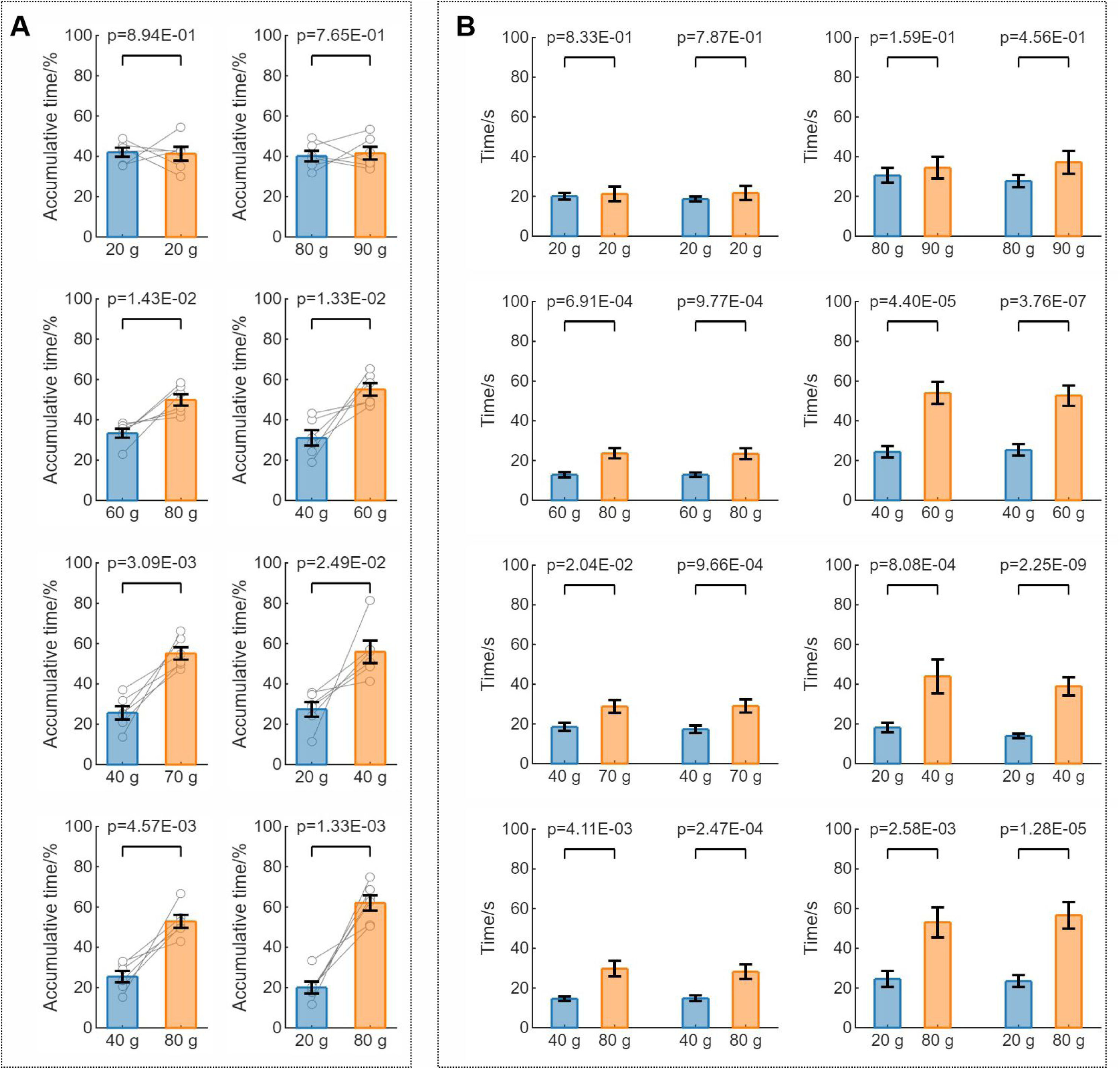
Quantity discrimination behavior during two-photon imaging experiments. (**A**) Quantity discrimination behavior for different amounts of food during the two-photon imaging experiments. Two-tailed Paired t-test; mean and SEM are shown; dots manifest data from individual mice (N=6 per group). (**B**) Time allocation in foraging zones during zone transitions when different amounts of food were presented. For given amounts of food (*e.g.*, 20 g versus 80 g), left two columns show transitions from more-amount to less-amount foraging zones, while right two columns show the reverse transition pattern. Two-tailed Wilcoxon signed-rank test; mean and SEM are shown.

**Fig. S8.**
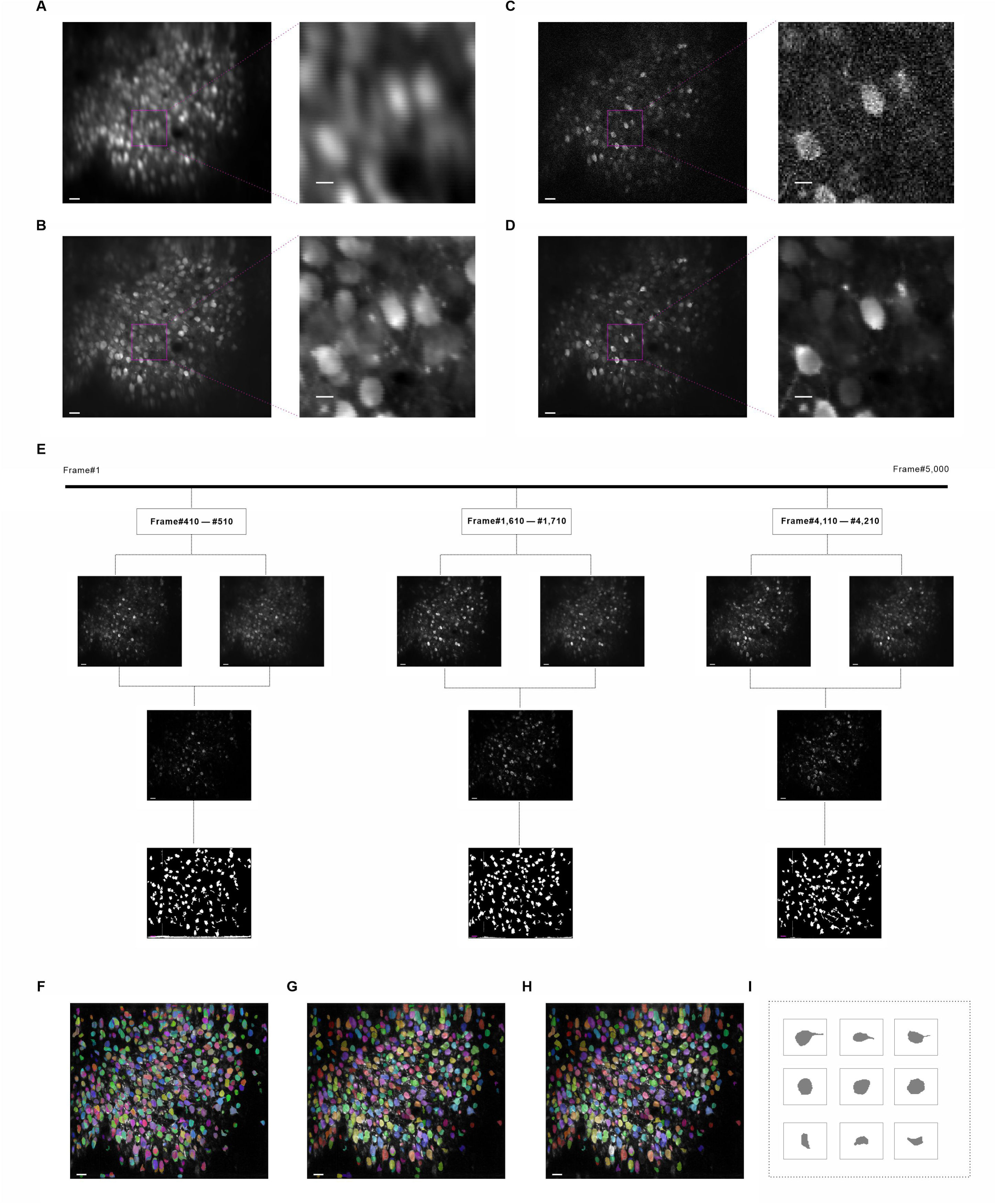
Two-photon imaging data processing. (**A**, **B**) Mean projection images of all frames before (**A**) and after (**B**) applying motion correction. (**C**, **D**) A single frame before (**C**) and after (**D**) noise reduction. (**E**) Three representative periodic blocks, each consisting of 100 consecutive frames. For each block, the maximum projection image (upper left), mean projection image (upper right), max-mean projection image (middle), and corresponding periodic mask (lower) are displayed. (**F**, **G**, **H**), Illustrations of the temporal mask (**F**), spatial mask (**G**), and soma mask (**H**). (**I**) Examples of typical soma. Scale bars: 30 pixels (**E**, left panels in **A**-**D**, **F**-**H**) or 10 pixels (right panels in **A**-**D**).

**Fig. S9.**
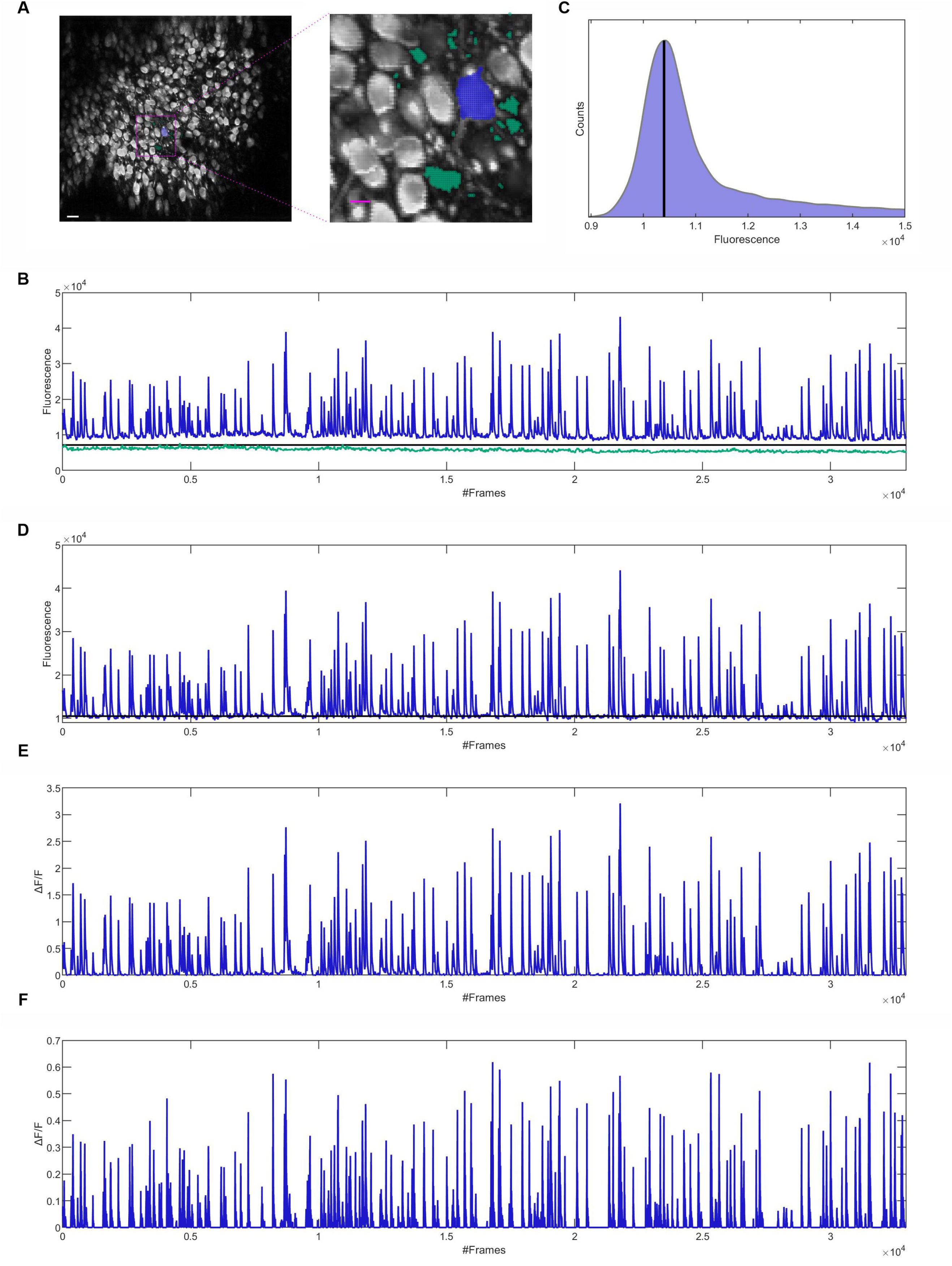
Extraction of neuronal activity signals. (**A**) A typical neuron (blue) and its corresponding neuropil (green). Scale bars: 30 pixels (left) or 10 pixels (right). (**B**) The average fluorescence intensity recorded from the neuron (blue) and its neuropil (green) throughout the entire recording session. The black line indicates the 3σ level of the background fluorescence. (**C**) Distribution of average fluorescence intensity across individual frames along with the baseline (black line). (**D**) The fluorescence trace of the neuron after correction for neuropil contamination and detrending to remove polynomial trends. The black line represents the baseline fluorescence. (**E**) The calcium signal of the neuron. (**F**) The event signal of the neuron.

**Fig. S10.**
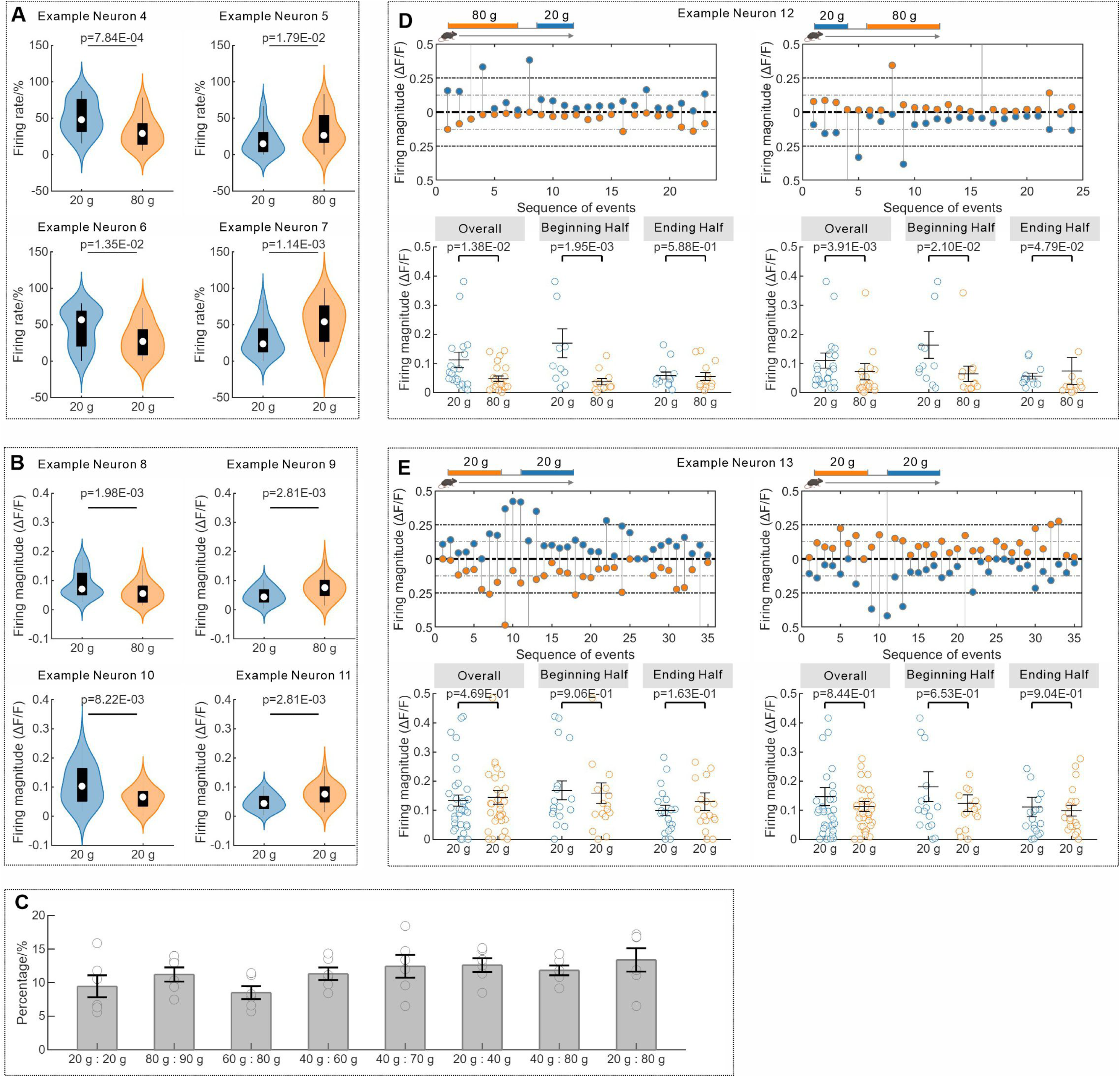
Neurons with preferential activities at a given time. (**A**, **B**) Typical neurons with preferential activities towards a specific amount of food, evaluated by comparing their firing rate (**A**) or firing magnitude (**B**) across different foraging zones. Two-tailed Mann-Whitney U test. (**C**) The percentage of neurons showing preferential activities for specific amounts of food. mean and SEM are shown; dots manifest data from individual mice (N=6 per group). (**D**, **E**) Preferential activities of two typical neurons in the PPC during the quantity discrimination assay for 20 g versus 80 g of food (**D**), or 20 g versus 20 g of food (**E**) at either the beginning or the ending half. Two-tailed Wilcoxon signed-rank test; mean and SEM are shown; dots manifest data from individual events.

**Fig. S11.**
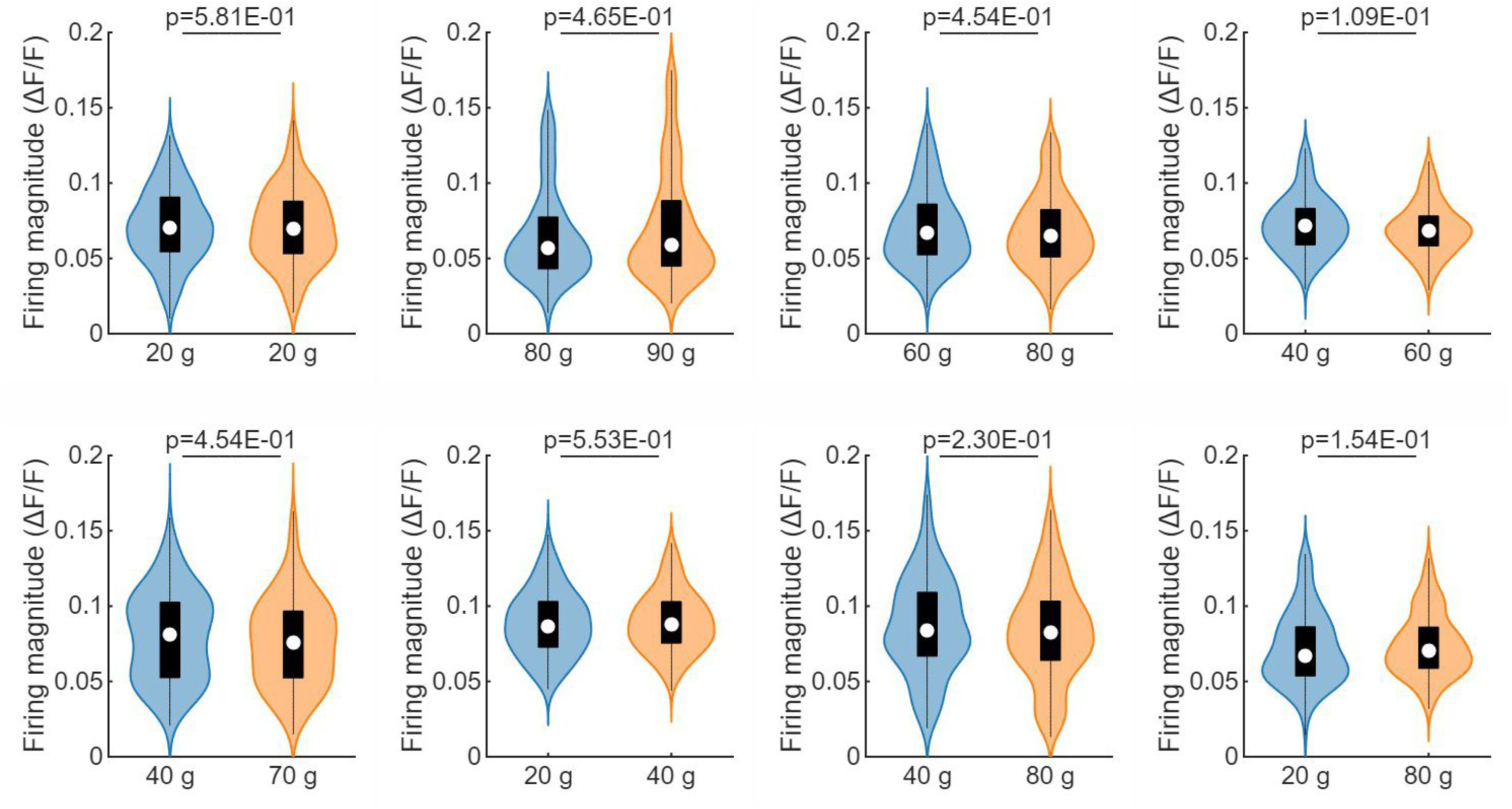
Firing magnitude of the PPC neuronal population. The firing magnitude of the PPC neuronal population when the animals were presented with different amounts of food. Two-tailed Mann-Whitney U test.

**Fig. S12.**
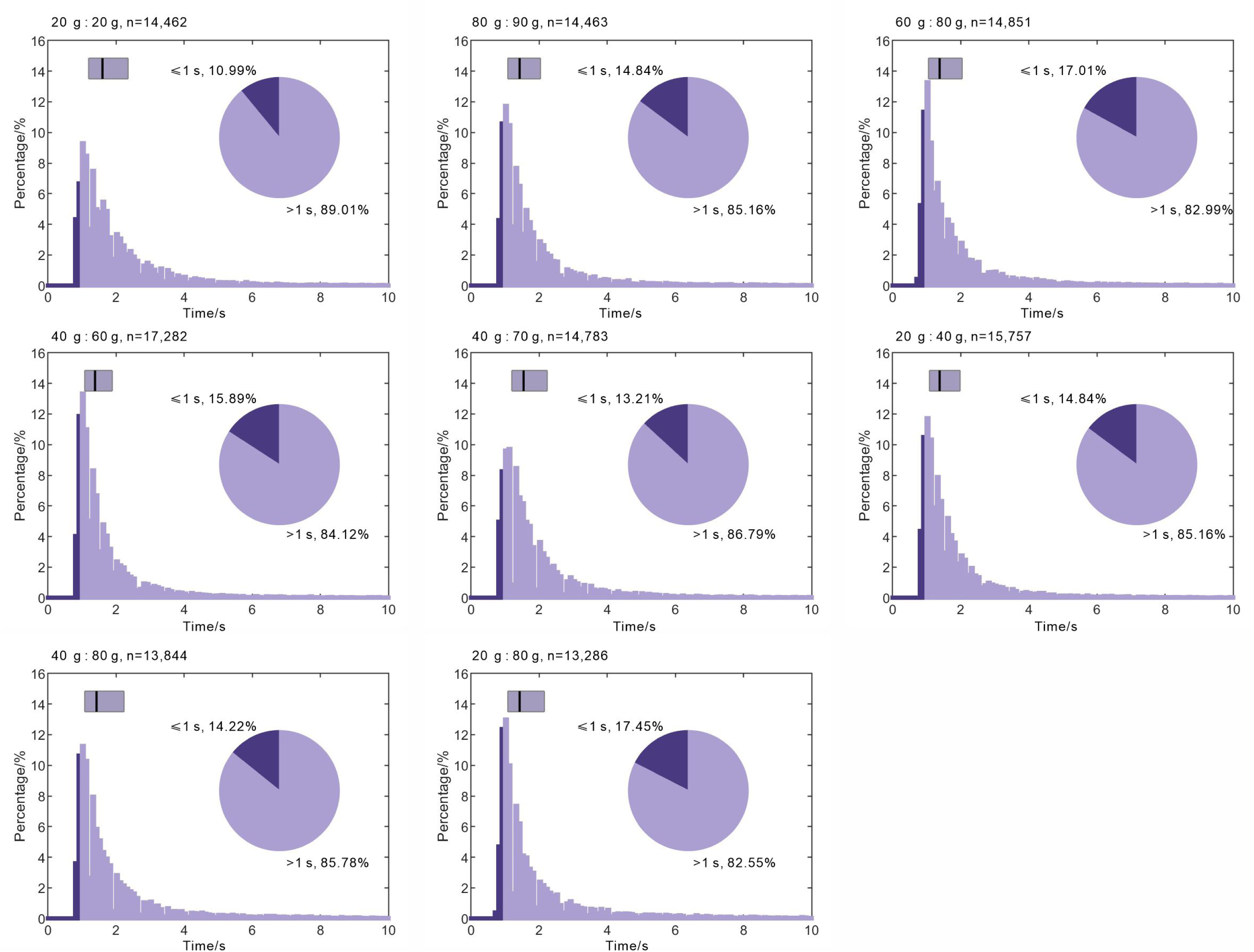
Duration of population synchrony. The duration of population synchrony in the PPC, illustrated with a pie chart depicting the proportion of patterns classified using a 1-second threshold. n indicates the number of population synchrony.

**Fig. S13.**
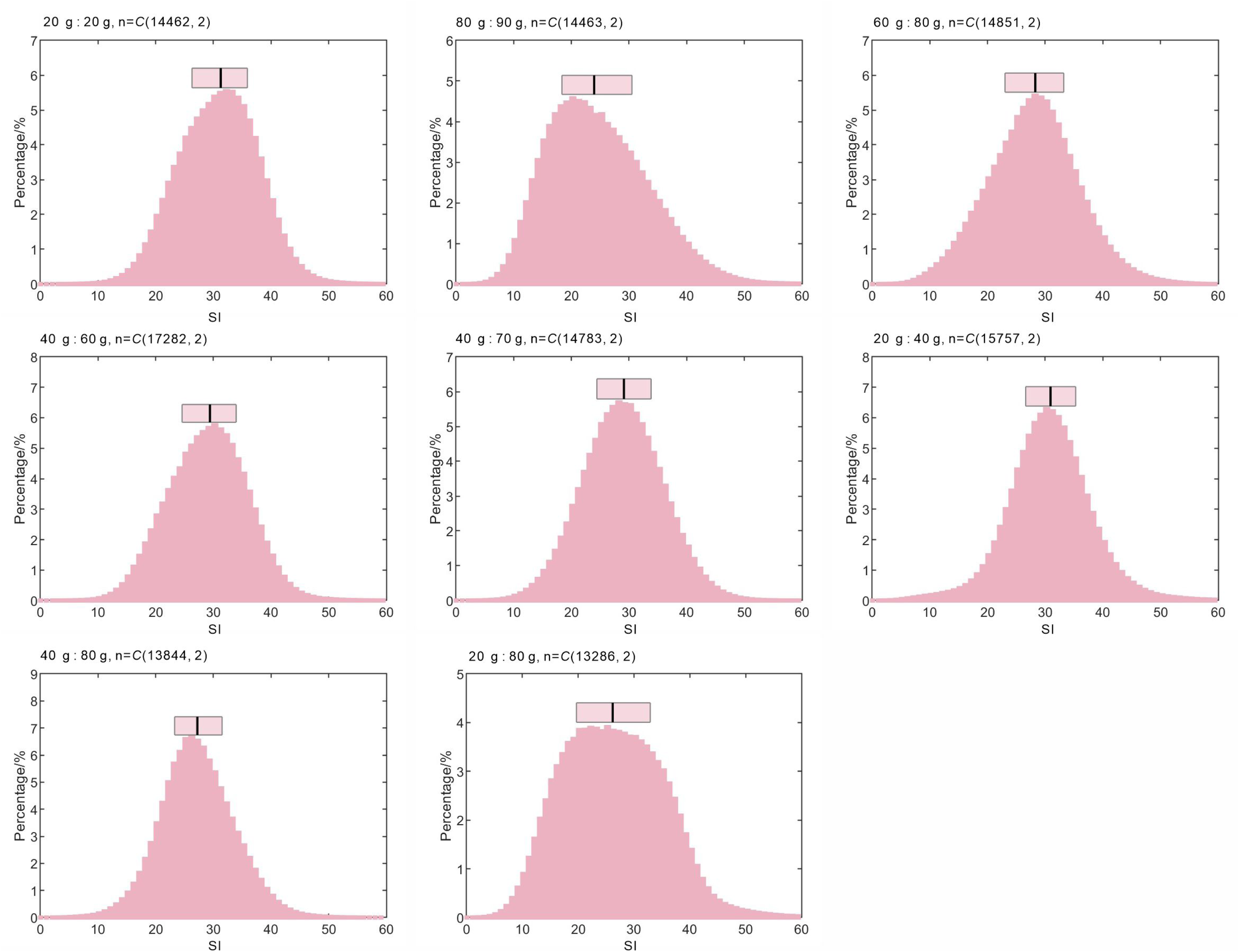
Distribution of similarity index. The distribution of similarity index (SI) between any pair of population synchrony. n indicates the number of population synchrony pairwise.

**Fig. S14.**
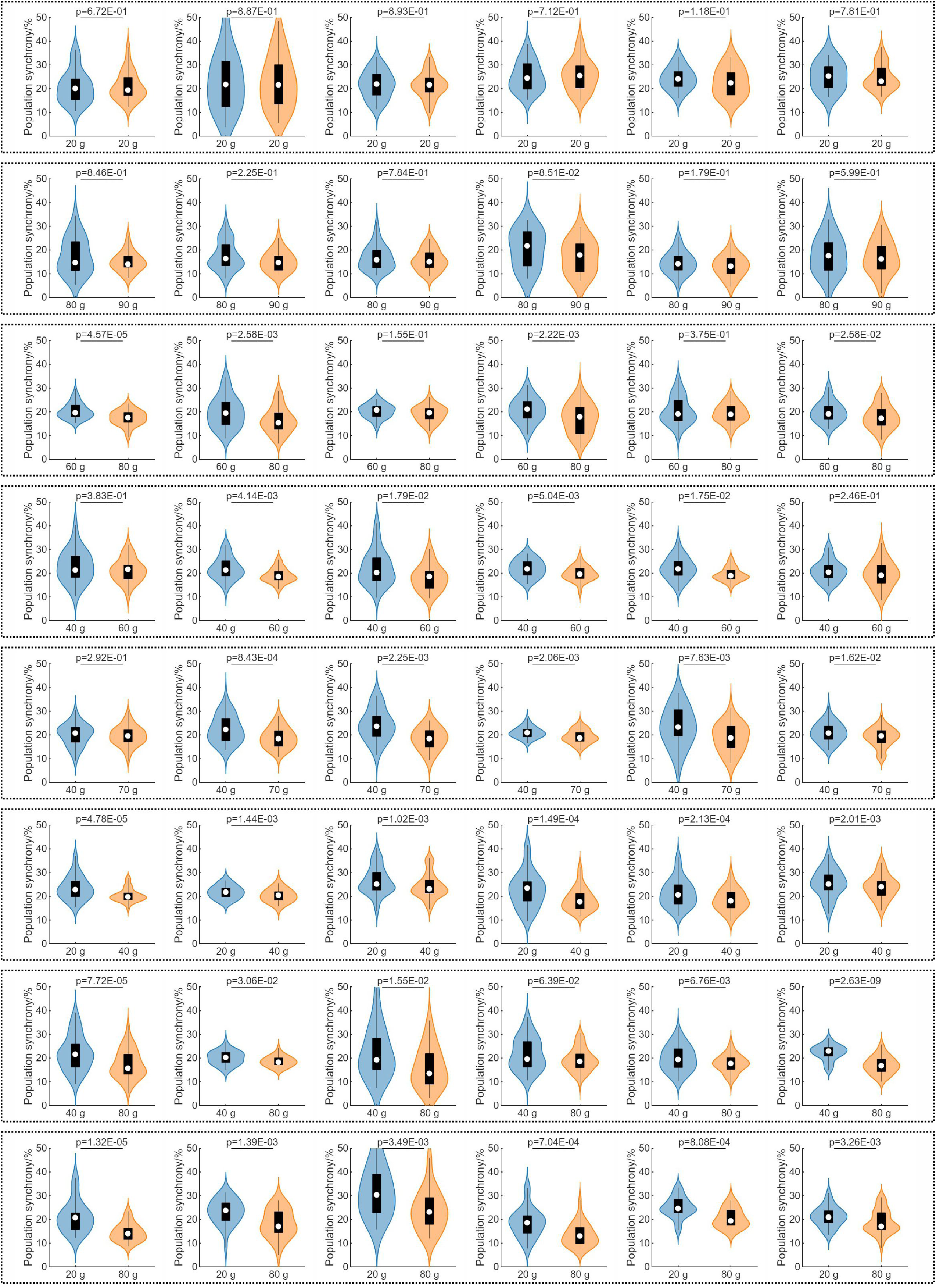
The levels of population synchrony during quantity discrimination. The levels of population synchrony in the PPC when individual mice were presented with different food amounts. Two-tailed Mann-Whitney U test.

**Fig. S15.**
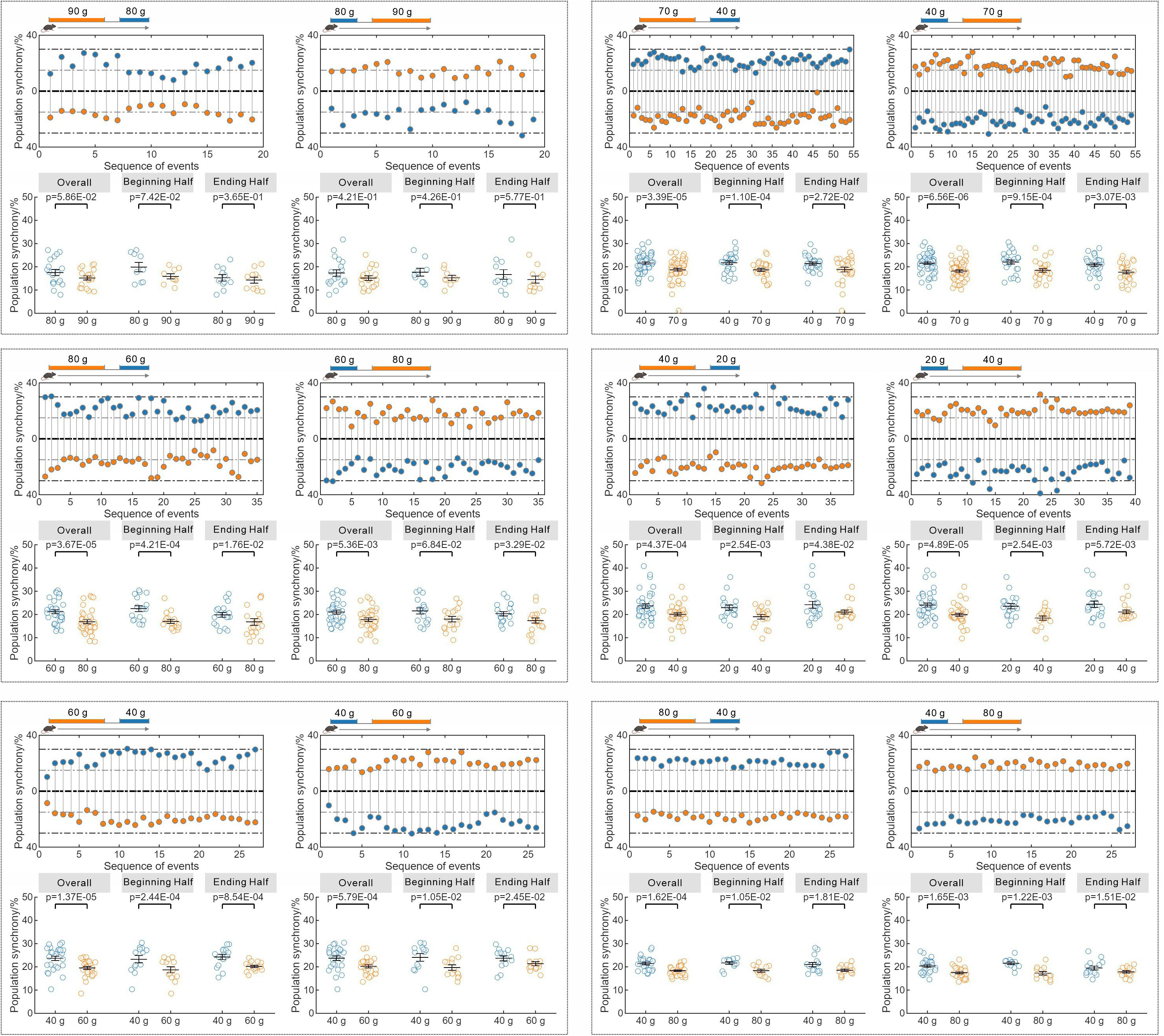
Changes in the levels of population synchrony during quantity discrimination. The levels of population synchrony in typical mice during the quantity discrimination assay for different amounts of food at either the beginning or the ending half. Two-tailed Wilcoxon signed-rank test; mean and SEM are shown; dots manifest data from individual events.

**Fig. S16.**
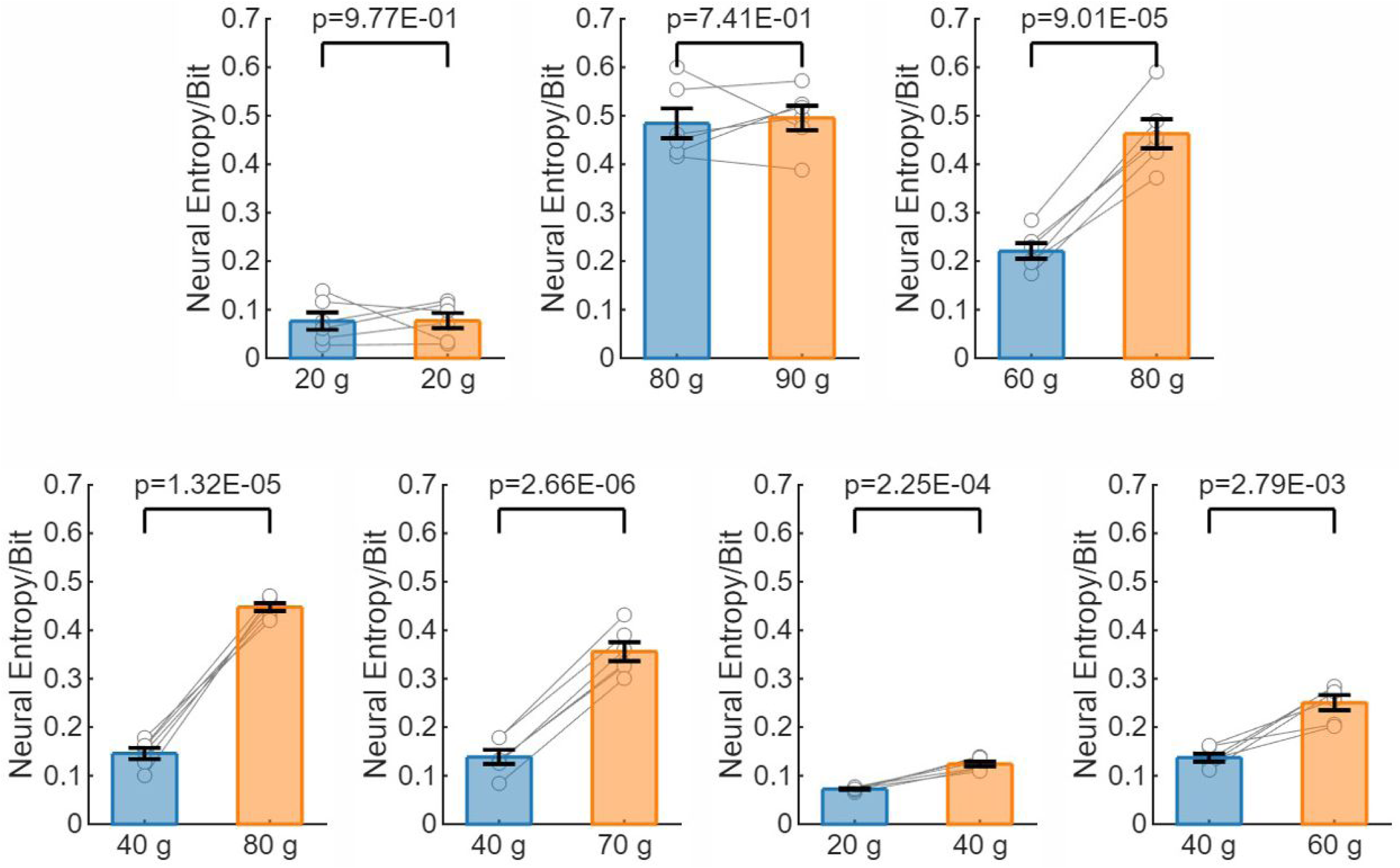
Neural Entropy values. The Neural Entropy values associated with different amounts of food. Two-tailed paired t-test; mean and SEM are shown; dots manifest data from individual mice (N=6 mice per group).

**Table S1.** The amounts of food and their ratios in the quantity discrimination assay.

| **Amounts of Food (g/g)** | **Ratio** |
| --- | --- |
| 40:40 | 1.00 |
| 80:90 | 1.13 |
| 50:60 | 1.20 |
| 30:40 | 1.33 |
| 20:30 | 1.50 |
| 40:70 | 1.75 |
| 40:80 | 2.00 |
| 20:80 | 4.00 |
| 15:90 | 6.00 |

**Table S2.** Two-photon imaging experiments.

| **Amounts of Food (g/g)** | **Number of Recorded Neurons** | **Duration of Population Synchrony (s) <sup>a</sup>** |
| --- | --- | --- |
| 20:20 | 1,002 | 2.03 ± 0.01 |
| 80:90 | 627 | 1.99 ± 0.02 |
| 60:80 | 898 | 1.88 ± 0.01 |
| 40:60 | 1,023 | 1.76 ± 0.01 |
| 40:70 | 791 | 1.99 ± 0.01 |
| 20:40 | 1,295 | 1.84 ± 0.01 |
| 40:80 | 935 | 2.04 ± 0.02 |
| 20:80 | 807 | 2.06 ± 0.02 |

<sup>a</sup> Values are presented as mean ± SEM (N=6 mice per group).

